# Data-driven approaches for genetic characterization of SARS-CoV-2 lineages

**DOI:** 10.1101/2021.09.28.462270

**Authors:** Fatima Mostefai, Isabel Gamache, Jessie Huang, Arnaud N’Guessan, Justin Pelletier, Ahmad Pesaranghader, David Hamelin, Carmen Lia Murall, Raphaël Poujol, Jean-Christophe Grenier, Martin Smith, Etienne Caron, Morgan Craig, Jesse Shapiro, Guy Wolf, Smita Krishnaswamy, Julie G. Hussin

## Abstract

The genome of the Severe Acute Respiratory Syndrome coronavirus 2 (SARS-CoV-2), the pathogen that causes coronavirus disease 2019 (COVID-19), has been sequenced at an unprecedented scale, leading to a tremendous amount of viral genome sequencing data. To understand the evolution of this virus in humans, and to assist in tracing infection pathways and designing preventive strategies, we present a set of computational tools that span phylogenomics, population genetics and machine learning approaches. To illustrate the utility of this toolbox, we detail an in depth analysis of the genetic diversity of SARS-CoV-2 in first year of the COVID-19 pandemic, using 329,854 high-quality consensus sequences published in the GISAID database during the pre-vaccination phase. We demonstrate that, compared to standard phylogenetic approaches, haplotype networks can be computed efficiently on much larger datasets, enabling real-time analyses. Furthermore, time series change of Tajima’s D provides a powerful metric of population expansion. Unsupervised learning techniques further highlight key steps in variant detection and facilitate the study of the role of this genomic variation in the context of SARS-CoV-2 infection, with Multiscale PHATE methodology identifying fine-scale structure in the SARS-CoV-2 genetic data that underlies the emergence of key lineages. The computational framework presented here is useful for real-time genomic surveillance of SARS-CoV-2 and could be applied to any pathogen that threatens the health of worldwide populations of humans and other organisms.

## Introduction

The Severe Acute Respiratory Syndrome coronavirus 2 (SARS-CoV-2) is a highly transmissible virus responsible for the current ongoing pandemic. SARS-CoV-2 is a positive-sense, single-stranded, 29,903 nucleotide long RNA genome. The virus’ genome is accumulating mutations at a steady pace since its introduction into human hosts. Genomic surveillance and the identification of variants of concern (VOC) and their impact on transmission, disease severity and immune response, are of tremendous importance to pandemic control, most notably in the context of worldwide vaccination efforts. In this context, an unparalleled wealth of chronologically and globally sampled viral genomes have been sequenced in a concerted public effort and submitted to public databases such as the Global Initiative for Sharing All Influenza Data (GISAID) (1).

In attempt to track the transmission and spread of emerging lineages, several lineage nomenclatures have been proposed, the most commonly used ones being the clades from NextStrain (phylogenetic approach) and Pangolin annotations (decision tree approach implemented in pangoLEARN) (2). Additionally, the World Health Organization assigns Greek letters to Variants of Concerns (VOCs), Variants Being Monitored (VBM), and Variants Of Interest (VOI). VOCs started emerging at the end of the first year of the pandemic, with the first one to be reported being the Alpha variant (3). Shortly after that, several other variants have attracted attention from the scientific community and public health bodies, amongst them Beta, Gamma, and Lambda, and more recently, the Delta variant which is becoming globally dominant (4).

Historically, phylogenetics is used to describe relationships between sequences, however, it is becoming more computationally intensive to use with the increasing number of closely related sequences made available on GISAID. Alternatively, by taking advantage of established population genetics paradigms that study the evolution of mutation frequencies in time, we may be able to describe and analyze the increasingly large datasets. For instance, an early study constructed a haplotype network to visualise circulating lineages of the SARS-CoV-2 virus (5). While phylogenetic approaches assume that the ancestral sequences are unobserved and represented by internal nodes, a haplotype network approach is appropriate when internal nodes actually are observed, because some sampled sequences are ancestral to others. This is the case for the current sampling scheme of pandemic sequences worldwide, despite many sampling biases (6; 7; 8). Tajima’s D (9), a classical population genetics approach, can be used to investigate SARS-CoV-2 lineage expansions (10). Dimensionality reduction techniques summarizing genetic diversity are also widely employed to investigate population structure. Principal component analysis (PCA) has been used to investigate the population structure of SARS-CoV-2 virus early in the pandemic (11). Uniform Manifold Approximation and Projection (UMAP), an unsupervised learning technique that has become increasingly popular in genomics in recent years, was previously used on GISAID sequences of SARS-CoV-2 (12). More recently, a novel unsupervised method, Multiscale PHATE (msPHATE) (13), was used for COVID-19 clinical data in combination with SARS-CoV-2 genetic data (14).

Here, we used genomics data collected in GISAID (1) from the fist year of the pandemic (January to December 2020, downloaded on January 19th 2021) to present a computational genomics pipeline for large-scale viral genetic profiling using a collection of established population genetics approaches and data visualisation strategies. Using these tools, we characterize the full scale of mutational diversity during the pre-vaccination phase of the pandemic. We show that these methods are useful to characterize the evolutionary steps undertaken by the virus during its early adaptation to the human host. This computational framework can help design efficient preventive strategies and derive fast response against viral adaptation to future therapeutic strategies.

## Results

### Viral Genetic Data Pre-processing

A key challenge in working with large datasets to extract meaningful information from genomic data is a careful pre-processing of the data to exclude low quality sequences, artefacts due to batch effects, and missing data. GISAID has very stringent submission guidelines and quality checks, which already guarantees data of good quality, but we’ve added several pre-processing steps (Step 1, Figure 1), which lead us to flag a series of systematic errors due to sequencing and bioinformatics methodologies (Methods). From the raw fasta file with 384,407 consensus sequences available in GISAID on January 19th 2021, we obtained a high-quality dataset of a total of 329,854 consensus sequences. Sequencing coverage across world regions is heavily biased, as we have 4194 sequences from Africa, 23499 sequences from Asia, 210624 sequences from Europe, 72774 from North America, 15009 sequences from Oceania and 3735 sequences from South America. We aligned these high-quality sequences and extracted all RNA substitutions compared to the reference sequence (NC 045512.2) (15), and imputed all sequences at 199 positions where derived allele frequency (DAF) was over 1% in at least one month using ImputeCoVNet (16). This novel approach is a 2D ResNet Autoencoder which has been shown to have an accuracy of > 99% and surpasses distance-based methods in terms of computation time (16), which is a major advantage in such a large dataset. We then built a harmonized database of RNA substitutions (Step 2, Figure 1) that contains a total of 24,802 mutated genomic positions (Table S1).

**Fig. 1:**
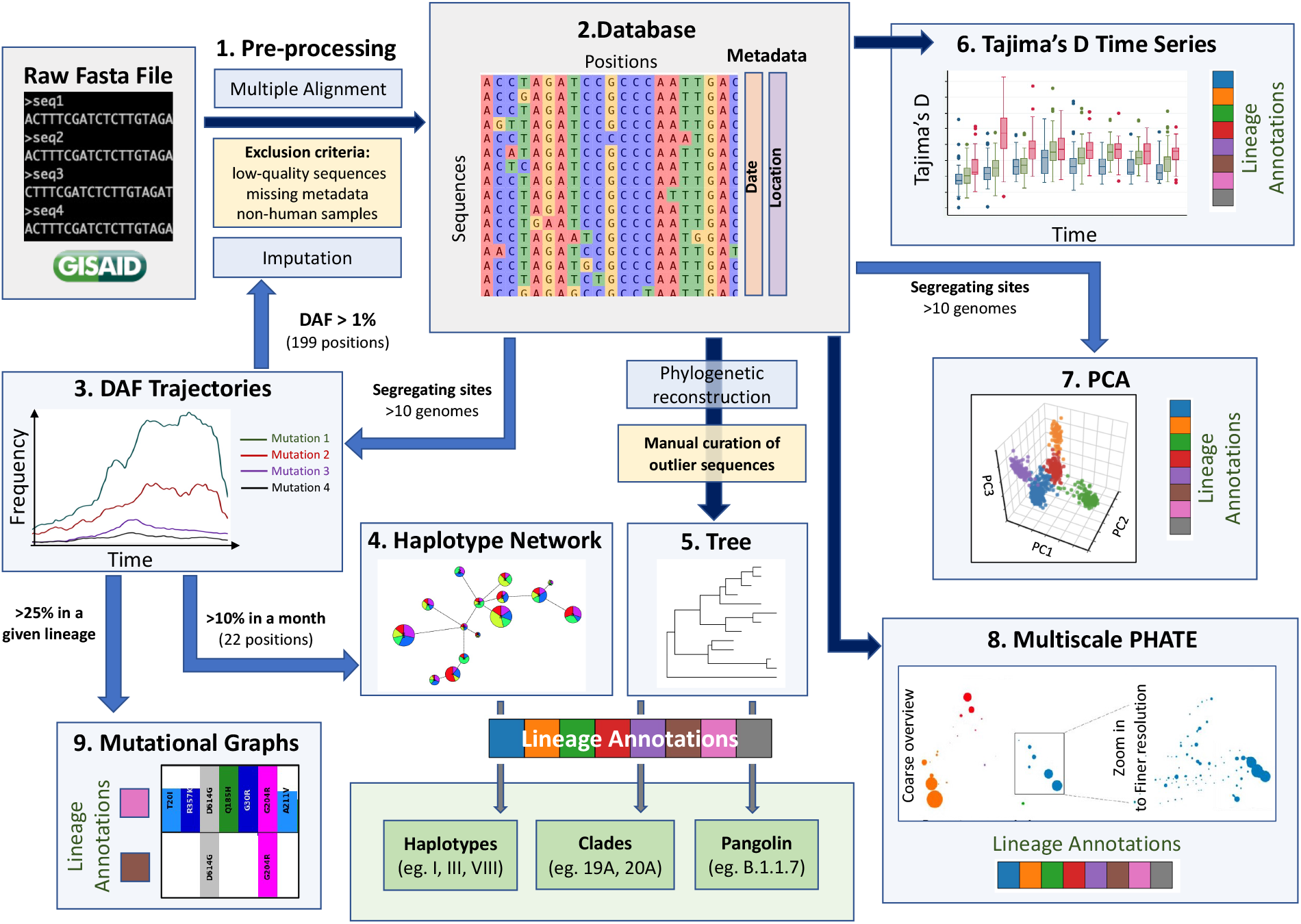
A Data-driven Methodological Pipeline for Analyzing Viral Genomic Data. This workflow recapitulates the major analysis steps used to analyse SARS-CoV-2 consensus sequence data inputted in GISAID during the first year of the COVID-19 pandemic. Dark blue arrows represent steps where all positions are kept (except spurious sites), blue arrows represents steps where subsets of positions are kept (indicated next to the arrow), yellow boxes represented filtering steps at the level of sequences, light bleu boxes represent the methodological steps and the main steps are numbered from Step 1 to Step 9. These population genetics and unsupervised learning methods constitute a comprehensive toolbox to allow the scientific community to efficiently monitor the evolution of the virus.

Since most of the consensus sequences in this dataset are from the UK (43%) and from USA (20%), we define two waves in the first year of the pandemic, corresponding to the two successive global increase of COVID-19 cases observed in countries that implemented strict containment measures in March 2020 and then relaxed these measures in the summer 2020, with the limitation that countries from the southern hemisphere had offset waves due to seasonality (17). The first wave encompasses sequences sampled from January to end of July 2020, and the second wave encompasses those sampled from August to December 2020 (18). We identified 20,403 substitutions within the first wave consensus sequences and 22,210 in the second wave, when compared to the reference genome (Table S1). Each of these mutated positions were further categorized based on their frequency of occurrence in the global host population during the first and second waves of the pandemic (Table S1). Singletons make up 23% and 17% of the first and second wave mutations, respectively: because these mutations are only seen in one sequence in each wave, they are most likely enriched in sequencing errors. Doubletons (mutations seen twice during a wave), account for 14% (first wave) and 11% (second wave) of the mutations, in line with an expanding population. There are 5% (first wave) and 11% of mutations found in more than 100 sequences in the first and second wave, respectively, where most impactful mutations on viral fitness will be found (19).

### Derived Allele Frequency (DAF) Trajectories Through Time

Expanding viral lineages will arbour a set of prevalent genetic mutations that will quickly increase in frequency over the course of a epidemic. To detect these mutations during the COVID-19 pandemic’s first year, we considered their derived allele frequencies (DAF) over time (Step 3, Figure 1). In the first wave, 20 RNA substitutions reached a DAF of 10% for at least one month (January to July) (Figure 2A). Four mutated positions are in linkage disequilibrium with each other, as evidenced by their DAF trajectories overlapping, meaning that they co-occurred together. Three of these substitutions are C-to-U mutations (C241U, C3037U, C4408U), the last one is A23403G in the Spike protein (S:D614G). These four substitutions have increased very quickly in frequency (Figure 2A, red DAF trajectory): this lineage was the first to become dominant and did so in a few months, climbing to 71% in March (Figure 2A). It is thought that S:D614G is the driver of this event and has been shown to have a selective advantage for SARS-CoV-2 which is conferred by an increase of transmission and viral load in the respiratory tract (20). Three consecutive co-occurring substitutions G28881A, G28882A, and G28883C also increased in frequency during the first wave (Figure 2A, purple DAF trajectory). It is the only change of three consecutive nucleotides, or triplet, that reached a DAF over 1% in the first year of the pandemic, an event that is unlikely to arise by chance alone, and could represent an adaptive change occurring on a codon. These three mutations span two amino acids in the N protein, leading to N:R203K and N:G204R. However, in the overlapping gene ORF9c (or ORF14) (21) they form a single codon mutated from GGG to AAC, causing a single missense change in the resulting protein (ORF9c:G50N). Interestingly, ORF9c is a novel gene in SARS-CoV-2 compared to known human coronaviruses (22), coding for a putative transmembrane protein. Additionally, this codon change significantly disrupt the RNA secondary structure of this specific region of SARS-CoV-2 genome, making a stable hairpin become a ‘wobbly’ loop (Figure S1). Finally, the first wave was marked by the increase in frequency of this group of five co-occuring substitutions: A1163T, T7540C, G16647T, C18555T, G22992A, G23401A (Figure 2A, yellow DAF trajectory). This lineage peaked in July 2020 and was mainly circulating and contained in Australia (23).

**Fig. 2:**
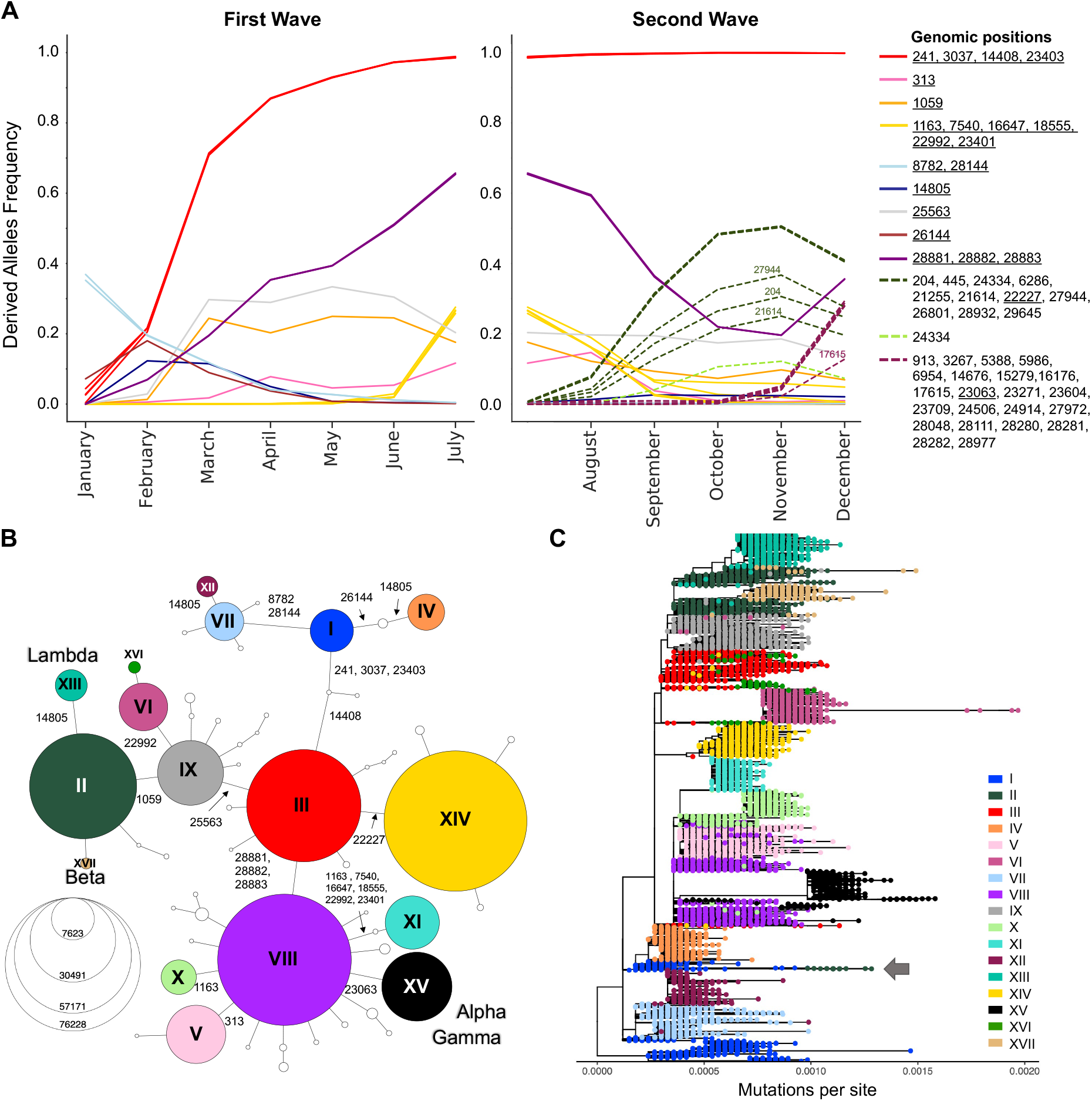
Viral Genetic Diversity During the First Year of the Pandemic. Derived Allele Frequencies (DAF) over time of representative high-frequency substitutions during the first (January-July) and second (August-December) wave of the pandemic. Only positions that exceed a DAF of 10% for a given month are shown. Positions with highly correlated DAF trajectories (*r*^2^ > 99) have the same line color. Solid lines are used for mutations appearing in the first wave. (B) Haplotype network representing genetic subtypes based on representative mutations (position underlined in A). Genomic positions that differs between two nodes (haplotypes) are specify on edges. Nodes are colored by haplotype and node size represent the number of consensus sequences for each haplotype. The 17 main haplotypes are annotated with roman numerals. (C) Divergence tree made from 15,690 SARS-CoV-2 consensus sequences using FastTree and TreeTime. The haplotype network built from prevalent mutations using all high-quality consensus sequences recapitulates well the phylogeny.

Overall, the second wave is marked by an increase in the number of highly transmitted mutations: from August to December, there is an increase in the number of substitutions with high DAF (Figure 2A), for a total of 53 mutations. However, among the 33 new high-frequency substitutions, the DAF trajectories show two well-defined groups, representing two different lineages: one with mutation S:A222V at genomic position 22227 (Figure 2A, dashed green DAF trajectories) and a lineage with mutation S:N501Y at position 23063 (Figure 2A, dashed plum DAF trajectories). The lineage with S:A222V corresponds to 20E in NextStrain annotations, and was first reported by Rambault and colleagues (24; 25). This lineage has accumulated 11 co-occurring mutations since its appearance, and was mostly seen in European countries. The lineage with S:501Y mainly corresponds to the NextStrain 20I, now commonly known as the Alpha variant, which accumulated 22 co-occurring high frequency mutations (Figure 2A) (26; 24; 25).

### Haplotype Networks for Fast Evolutionary Clustering of Sequences

The generation of haplotype networks is a widely used approach for analysing and visualizing the relationships between sequences within a population (27; 28). The nodes of the network are haplotypes, edges represent mutated genetic positions that vary between two nodes (Figure 2B). The size of the nodes generally varies to represent the number of sequences for a specific haplotype. In the case where there is very little to no recombination, this approach results in a minimum spanning tree. To keep the number of haplotypes tractable for informative visualization, we defined them by selecting the 22 mutations displayed in the DAF trajectories that are most representative (Methods, Figure 2A underlined positions, Table S2). We generated a haplotype network (Step 4, Figure 1) using a technique that takes the time of sampling into account (Methods). We included the 122 haplotypes with more than 10 sequences in our representation, ignoring rare events. The final haplotype network includes 17 main haplotypes (Figure 2B), representing the main genetic lineages circulating during the pandemic’s first year, and several “descendant” haplotypes. Haplotype I include all sequences with ancestral states at each position (ie. reference haplotype), and haplotype XV corresponds mainly to the Alpha variant, and differs from the reference haplotype at 8 positions.

To further visualise the genetic diversity of the virus in the first two waves, we looked at the viral diversity during each wave of the pandemic using the haplotype network (Figure 3A,B). During the first wave, the haplotype network shows the presence of nine haplotypes with more than 200,000 sequences (II-V, VII-IX, XI, XII) diverging from the ancestral haplotype I (Figure 3A), with haplotypes II, III and VIII being the most prevalent, all carrying the S:D614G mutation. Most haplotypes are seen in several regions of the world, albeit at different frequencies (eg. V arose mainly in Asia and II mainly in North America), except for haplotype XI. This latter lineage (Pangolin D.2) was mostly circulating in Australia (encompasses 92.8% of all high-quality Australian sequences in GISAID in July 2020). First detected in June 2020, this variant almost completely vanished as of October 2020. The second wave is marked by a critical decrease in the prevalence of haplotypes without the S:D614G mutation (I, IV, VII and XII) that have almost become extinct by August 2020, and the fast increase in prevalence of haplotypes VI, XIV and XV, arising mainly in Europe. Other novel region-specific haplotypes (XIII in North America, XVI in Europe, XVII in Africa) arose during this wave (Figure 3C). Additionally, the second wave was marked by the appearance of lineages with mutational jumps on haplotype VI and XV (Supplementary Text, Figure S5) using root-to-tip estimates of mutational rates in each haplotype lineage (Methods).

**Fig. 3:**
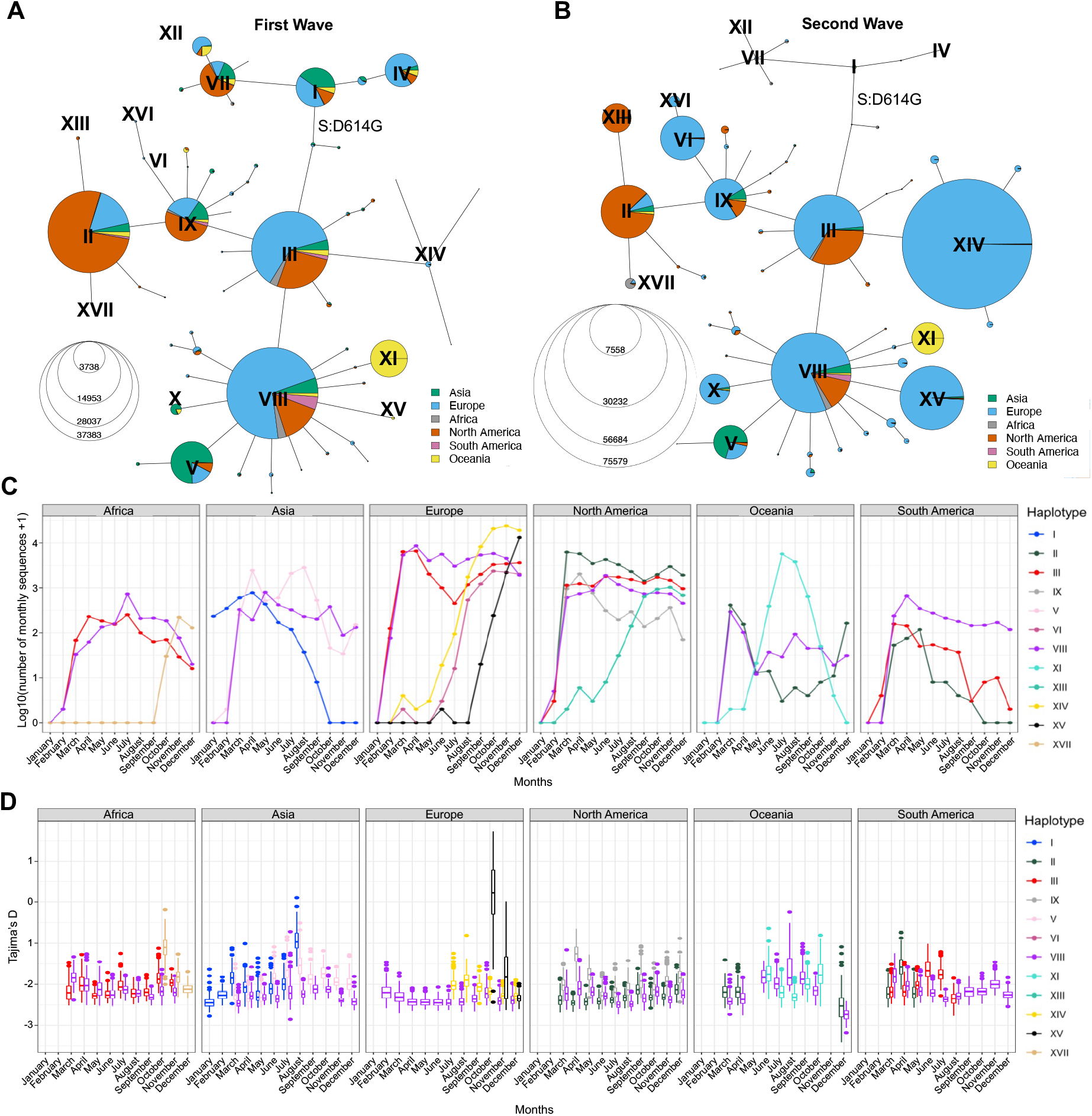
Distribution of SARS-CoV-2 Sequences in Space and Time. Haplotype network of the first (A) and second (B) wave. Node size represent the number of consensus sequences for each haplotype and pie charts represent continental proportions for each haplotype. (C) GISAID consensus sequence counts (on a log10 scale) of the most prevalent haplotypes in each continent during the first year of the pandemic. (D) Tajima’s D estimates of the three most prevalent haplotypes in each continent for the first year of the pandemic. Boxplots represent 500 resampled estimates of Tajima’s D from random resamplings of 20 genome sequences for each month with at least 20 sequences. Both the haplotype network and Tajima’s D are insightful tools to detect expanding lineages at a given point in time.

### Comparison to Phylogenetic Reconstruction

To compare the haplotype network to a more standard phylogenetic approach (Step 5, Figure 1), we generated a divergence tree using FastTree along with other complementary tools (Methods), as recommended in multiple published pipelines (29; 30). With these approaches, using all 329,854 sequences is unfeasible, therefore sub-sampling is needed (Methods) which can bias the representation of the circulating diversity of the viral population. In contrast, the haplotype network can be built with all available sequences, while recapitulating the phylogenetic structure quite accurately, despite some lineage splitting, particularly of haplotype I (Figure 2C). The phylogenetic approach sometimes wrongly combines very distant lineages, for example lineage I and II (Figure 2C, arrow) where further inspection of these sequences put into doubt the relationship reported by the phylogenetic tree. Indeed, the haplotype annotations allow to identify problems in the phylogeny that would otherwise go undiscovered, technical issues that can then be corrected by additional tools and manual adjustments (Supplementary Text, Figure S3). Additionally, the haplotype network allows easy representation of recurrent mutations, for instance, the C-to-U mutation at position 14805 (Figure S4A), which emerged on three different backgrounds (IV,XII,XIII).

### Comparison to Other Lineage Annotation Systems

Our haplotype definition is well in line with NextStrain lineage definition (Figure S2A), though some distinctions exist. For instance the haplotype approach can differentiate sequences from haplotype I and IV, differing by two substitutions (genomic positions 26144,14805), and from III and IX, differing at position 25583, whereas NextStrain does not differentiate these lineages, grouping them into 19A and 20A, respectively. In contrast, our categorization of sequences by haplotype and the Pangolin annotation methodology (3) do not concord well (Figure S2B), with, for instance, the B.1 lineage spanning many genetically distant haplotypes: sequences from haplotype II and VIII can be assigned to the same generic B.1, while these differ by a total of five high frequency mutations, including the triplet at 28881-28883. Conversely, lineages B.1.1, B.1.2, B.1.5 are sublineages of haplotypes VIII, II, and III, respectively, defined by mutations that never surpassed 10% worldwide. These inconsistencies between the genetic background of different Pangolin lineages, and the greater granularity observed compared to NextStrain lineages justifies the usage of haplotype categories for the analyses of genetic evolution of SARS-CoV-2. However, the haplotype definition based only on these 22 most prevalent mutations in 2020 does not differentiate all VOCs. For example, the sequences of the Gamma variant discovered in Brazil (31), which also has the substitution at position 23063 (S:N501Y) as well as the triplet, are grouped with Alpha sequences in haplotype XV.

### Time Series Change of Tajima’s D statistic to Detect Lineage Expansions in Real Time

The haplotype network representation informs on the overall size of clusters but lacks information on the lineages’ expansions over time, especially by geographical region. Population genetic statistics, most notably Tajima’s D, has been used to estimate epidemiological parameters of pandemic influenza A (H1N1) (32) and can detect population expansion and contraction events. Specifically, a strongly negative Tajima’s D value indicates a rapid population expansion, resulting in an excess of low-frequency alleles in the population. However, Tajima’s D is very sensitive to population structure and is therefore not meaningful when applied on a global scale. The genetically-informed grouping of sequences by haplotypes as well as stratification by world regions, however, allows to get closer to well-mixed viral populations, where Tajima’s D can be applied. For prevalent haplotypes in each continent, we thus computed Tajima’s D for each month of 2020 (Step 6, Figure 1) while controlling for sample size (Methods). We recognize that the uneven sequencing coverage across the continents may bias mutation rate estimates, therefore we avoided comparing regions. We then correlated Tajima’s D to the number of sequences per haplotype per continent and observed a moderate negative correlation (mean adjusted R2 = 0.24, s.d. = 0.23, Figure S6A). This correlation is significant enough such that the time series of Tajima’s D recapitulates the major variations in the number of sequences per haplotype per month across each continent (Figures 3C and 3D). Indeed, we observe a decrease of Tajima’s D through time for several haplotypes that either took over in a specific region or are known to have become dominant (Figures 3C,D and S6B,C). For instance, in Africa, haplotype XVII (Beta) appears in October 2020 with a high Tajima’s D value compared to dominant haplotypes VIII and III and then shows a fast decrease of Tajima’s D in the next months, consistent with population expansion (Figure 3D). This correlates with the drastic increase in number of sequences seen at the end of 2020 (Figure 3C), reflecting the emergence of Beta in South Africa (33). Another striking event of this type is seen in Europe where the rapid increase of XV (Alpha) coincides with a steep decrease in Tajima’s D from October to December (Figure 3C,D and S6B). These events are also consistent with observations from Singh and Yi (2021) who tracked the spread of the corresponding Nextstrain clades (XV: Nextstrain 20I; XVII: Nextstrain 20H; (34)).

In North America, Tajima’s D for haplotype IX also shows a marked decrease from April to June, suggesting an expansion of this lineage, although its prevalence in the sampled population from GISAID did not increase (Figure 3C). This may reflect undersampling of specific populations in North America. Conversely, the rapid rise of haplotype XIII (Lambda) is captured both by Tajima’s D and sequence counts (Figure S6C). Other types of events can be seen, such as loss of lineages and lineages causing outbreaks that are quickly contained. An example of the former is seen in Asia where SARS-CoV-2 emerged (35), where Tajima’s D for haplotype I sequences increases with time, suggesting a contraction of population size of the ancestral lineage, with haplotype VIII and V becoming dominant haplotypes by the end of 2020. The signature of a contained outbreak is seen in Oceania, where Tajima’s D distribution across time is U-shaped, indicating an increase in population size followed by a contraction, in line with sequence counts (Figure 3C). In South America, sample counts per haplotype suggests that haplotype VIII took over II and III by the end of 2020, but the low Tajima’s D values in August 2020 are inconsistent with a decrease of these lineages and suggests that the number of sequences assigned to these haplotypes is an underestimate in that region.

### Visualizing viral fine-scale population structure using Principal Component Analyses (PCA)

To detect and visualize fine-scale structure in genetic variation, a common statistical approach is to use PCA of coded alleles at mutated positions segregating in populations. The projection of sequences onto the principal components is known to reflect the underlying (generally unknown) genealogical relationships between haploid sequences (36). In the viral populations of SARS-CoV-2 from 2020, we performed PCA (Step 7, Figure 1) on viral mutations present in at least 10 sequences from the first and second wave (Figure 4). The first two PCs describe the most variation between sequences and clearly show discrimination between haplotypes dominating during either of the two waves (Figure 4A,B). The coordinates of clusters relative to one another in the first wave concord with the haplotype network representation (Figure 3A): on PC1, the haplotypes without S:D614G are separated away from VIII and XI carrying the triplet mutations, whereas PC2 separates them from haplotypes II and IX, with haplotype III located at the center, in line with its intermediate position in the haplotype network. Additionally, the distance in the PC1/2 space between haplotype groups appears to recapitulates very well the genetic distance between them. For instance, haplotype III sequences are at least four mutations away from I, three mutations away from VIII, and two mutations away from II, reflecting distances between groups on the graph (Figure 4A). In the second wave, PC1 and PC2 do not reflect these phylogenetic relationships as much, but rather highlight the most divergent groups, the major second wave lineages (Figure 4B). Both PCs show the XV group as an extreme group, which is explained by the major mutational jump of 22 mutations from haplotype VIII background defining Alpha. We can however see a subset of XV sequences clustering with VIII sequences, which are either precursor sequences of Alpha, or Gamma sequences. Beyond PC1 and PC2, other PCs from the two waves show additional structure within and between haplotype subgroups (Figure S7). For instance, PC3/4 of the first wave sequences show the divergence of haplotype XI (Figure 4C), the lineage dominating in Australia in the summer, whereas PC4/5 of the second wave sequences show the emergence of the haplotype XIII lineage (Beta) from its ancestral background on haplotype II (Figure 4D).

**Fig. 4:**
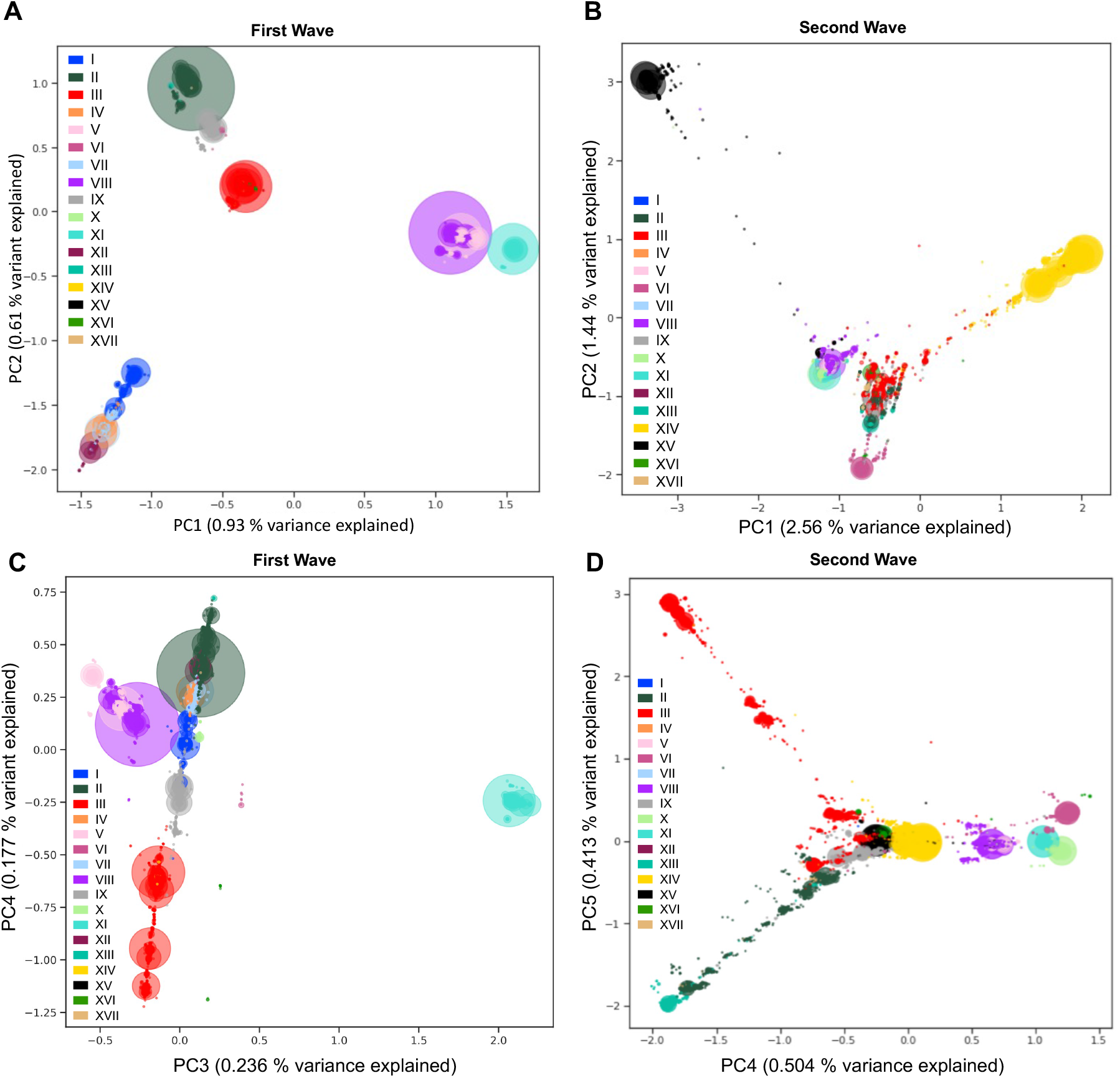
Viral Population Structure During the First and Second Wave of the Pandemic. Principal Component Analyses (PCA) of genetic diversity of the first (A,C) and second (B,D) wave consensus sequences reveal the population structure of the 17 main haplotypes in each wave. Genetic variation present in at least 10 genomes is used. The PCA is computed with all sequences, and only the sequences from the 17 main haplotypes are projected. Identical sequences are projected onto the same coordinates, therefore the number of sequences represented by each point is proportional to the size of the dots, with added transparency. PC1 and PC2 show differentiation between the main lineages from the two waves (A,B). The variant responsible for the Australian outbeak stands out clearly on PC3 from the first wave (C) and the Lambda variant sequences (XIII) are shown as the most distal subgroup on PC4 and PC5 from the second wave (D), in opposition to sequences from haplotype VI (on PC4) and subgroups of haplotype III (on PC5). The PCA recapitulates insightful characteristics of the evolutionary relationships of sequences and identifies major lineages from the two pandemic waves.

We then used UMAP to further summarize the 20 first PCs, explaining a total of 2.82% and 7.32% of genetic variance in the first and second wave respectively, in a 2-dimensional plot (Methods). UMAP has shown some potential to describe population structure from genetic variation (37), but in this case, despite the UMAP representation showing some clustering by haplotypes, many small groups of sequences are scattered around in an unstructured way (Figure S8), indicating that this technique may not be the most appropriate for visualisation of genetic diversity in SARS-CoV-2 genomes, even when exploring several parameters (Methods).

### Multiscale PHATE (msPHATE) for Data-driven Viral Lineage Groupings Across Granularities

PCA reveals insightful characteristics of the genetic data, and identifies major lineages, but grouping of data points (here, sequences) in PCA derived from genetic variation is known to be heavily influenced by uneven sampling of sequences, which means that the number of sequences sampled within sub-lineages will influence distances between subgroups (36). This is not a desirable attribute for early detection of new differentiated sub-lineages, which are often sampled in lower numbers compared to the other well-established lineages. We thus explored the use of msPHATE on SARS-CoV-2 genetic data, a novel unsupervised learning technique that showed promising results on biological data (13), to enable the identification of additional structure within haplotype groups using whole viral consensus sequences (Step 8, Figure 1). The method creates a tree of data granularities that can be cut at coarse levels for high-level summarizations of data, or at fine levels for detailed representations on subsets. The coarse level representation clusters sequences within groups which recapitulate the major circulating viral haplotypes during each wave (Figure 5). Across different granularities, msPHATE is able to uncover viral sub-lineages: a finer level representation of the first wave sequence data reveals three separate clusters of haplotype II, and one of haplotype III clusters splits into five sub-clusters (Figure 5A, right panel) and each subgroup mutational landscape can be inspected (Supplemental Text, Figure S9B-D). One of the five subgroups is made of sequences belonging to haplotype XIV, which only became particularly prevalent during the second wave in Europe. This shows the potential of msPHATE to identify emerging lineages.

**Fig. 5:**
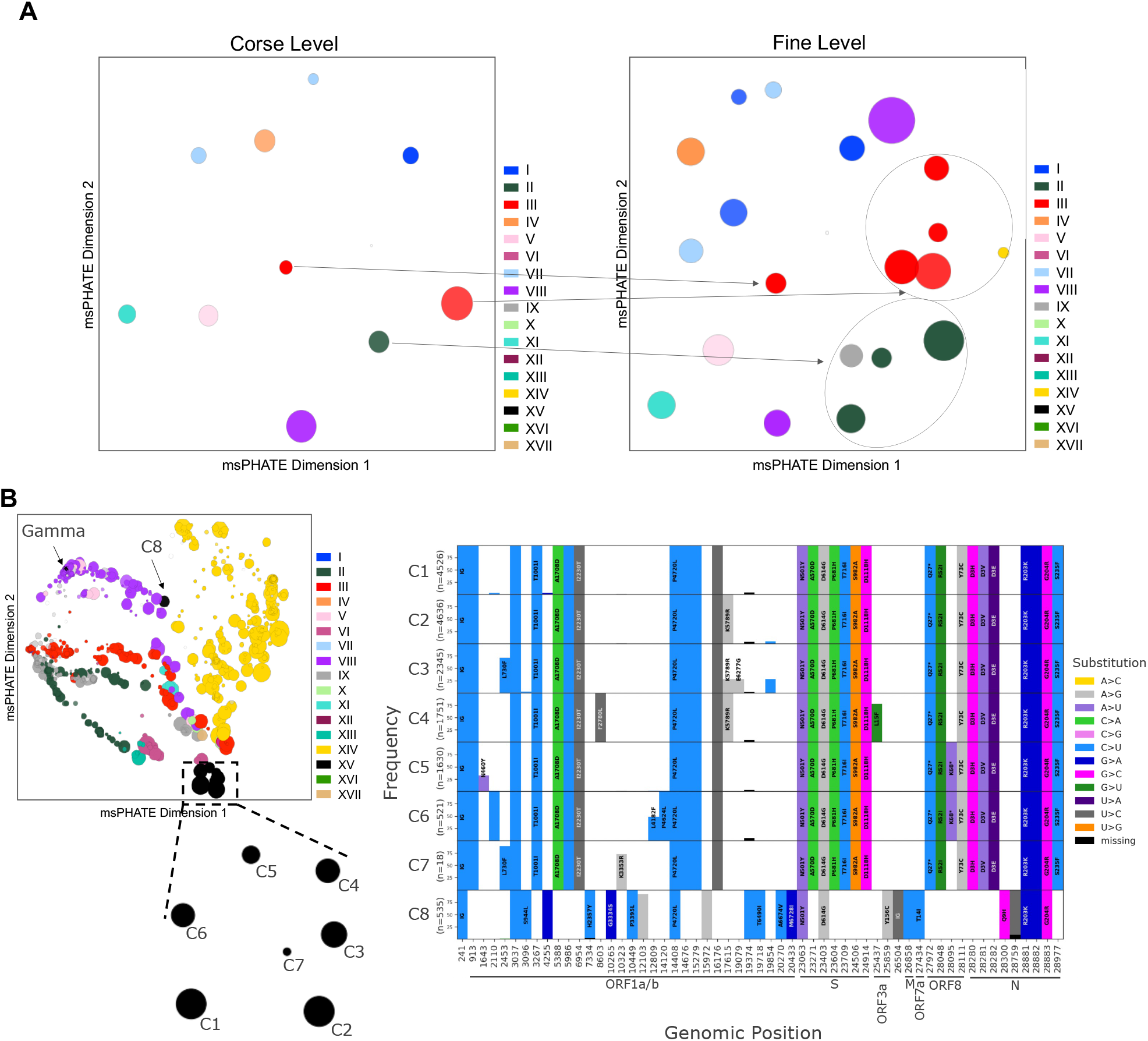
Viral Population Structure Using a Novel Reduction Technique Multiscale PHATE (msPHATE) (A) Coarse (left) and fine (right) grain visualizations of msPHATE embedding generated from genetic diversity of the first wave consensus sequences. The clusters are colored by majority vote. Subsets of haplotype II and III can be seen in the finer level. (B) Fine grain visualizations of msPHATE embedding of genetic diversity of the second wave consensus sequences (left). Zooming into haplotype XV clusters reveals 8 clusters (sub-lineages). Gamma variant is part of an VIII cluster that is marked with a star symbol. Mutated positions in each cluster group of haplotype XV are listed along the x-axis in the order they appear on the genome (right). Mutations in at least 25% of the samples for each haplotype XV cluster are represented as a bar, and the corresponding amino acid changes are shown on the bars (blank bars represent synonymous mutations). Genomic position annotation was done using SnpEFF (38). All the clusters share the mutation S:N501Y which characterize the haplotype XV. msPHATE embeddings allow the early detection of small subgroups of emerging lineages and can highlight putative variants of concern.

The finer level of the second wave embeddings shows very distinct patterns for the main circulating sub-lineages. There were many different sublineages for XVI sequences compared to XV subgroups (Figure 5B, left panel). Zooming into the haplotype XV clusters, we identified several sub-lineages (C1-8), which showed different mutational landscapes. For each of these subgroups, we can detailed their mutational landscape by deriving mutational graphs (Step 9, Figure 1). These graphs are stratified histograms of DAF for genomic positions that differ from the reference sequence in at least one subgroup, allowing comparison of lineages’ mutational landscape. The mutational graph of the eight XV subgroups, as defined by msPHATE, shows unique mutational events that define specific clusters (Figure 5B right). The C1 cluster shows all Alpha-defining mutations, but no other mutation has reached a DAF of 25% within this group, whereas the other clusters C2-7 have accumulated additional mutations. Notably, C5 and C6 present the ORF8:K68* nonsense mutation at genomic position 28095, a mutation at low frequency by the end of December 2020, but which quickly increased in frequency in the first months of 2021. Indeed, it was found in over 80% of Alpha sequences by September 2021, revealing a potentially beneficial mutation on Alpha background (Figure S10). While Gamma was found in a msPHATE grouping with other VIII sequences (Figure 5B), we detected an additional subgroup of XV, cluster C8, which does not have the same mutational landscape as the rest of the XV sub-lineages. This lineage (Pangolin lineage AP.1) (19), despite having experienced an independent mutational jump of 7 mutations at once, disappeared early in 2021.

## Discussion

The world-wide efforts to sequence and share thousands of viral genome sequences made possible in depth tracking of the evolution of the SARS-CoV-2 over time as it spread across the world. However, processing, analysis, and interpretation of hundreds thousands of sequences and mutation events is a challenging task (12). Here, we first propose a pre-processing pipeline to improve downstream analysis by using only high-quality data. Imputation of missing alleles at key positions facilitated haplotype annotation and therefore lineage tracing. Detection of spurious sites due to sequencing errors and bias is also an important step, and data analysts should be careful about these when processing large amount of genomic data, especially in a small genome of 29Kb, where by now, every position in the genome has been affected by sequencing errors.

We tested population genetics approaches to explore and identify emerging SARS-CoV-2 lineages. Allele frequency tracing through time is a widely used population genetics approach that can monitor circulating lineages over time. SARS-CoV-2 evolution experienced a period of relative evolutionary stasis during its first months infecting human hosts, which is consistent with other reports (19; 39). Mutations with increased fitness started emerging and led to a number of variants of concern, some of which are the leading cause of current infections. Indeed, the second wave of the pandemic was marked by the appearance of lineages with mutational jumps, explaining the increase in the number of substitutions, which may reflect adaptive steps in the evolutionary trajectories of SARS-CoV-2 within the human host, with the virus acquiring selective advantages to the host immune system (40; 41; 42; 43). Finally, this approach of analyzing worldwide frequencies of mutated positions allows a comprehensive picture of the virus’ strategies to developing new mutations and therefore lineages, exemplified by the observation of the Alpha variant emerging, starting in October (Figure 2B) but was only reported at the end of December (3).

The nomenclature has become increasingly complex with the number of lineages and sub-lineages emerging throughout the world. We explored tracing of lineages using a haplotype network, that was generated using the most frequent mutational events in a given time period, to clarify the genetic background of VOCs in 2020. Interestingly, Delta variant is accompanied with one of the triplet mutations, G to A at position 28881 (N:R203M), such that this variant emerges in the third wave network. It is easy to accommodate new mutations as new haplotypes can be generated with any mutational events of interest on any region of interest, such as the Spike protein, the main vaccines target currently.

We captured the expansion and decline of major circulating haplotype populations in each continent, and correlate them to sequences counts using Tajima’s D, a classical neutrality test statistic. Some inconsistencies between Tajima’s D predicted expansions and decrease in sequence counts could be an indication of undersampling in a given region. Sampling biases are numerous in this dataset, and attempts to correct them may also lead to other systematic errors. Furthermore, it is a very unbalanced dataset in terms of sampling countries, with 144,376 (44%) of the sequencing effort during the first year done in the UK. Global strategies aiming at a more uniformly distributed between countries sequencing effort would be able to identify earlier emerging variants. Population structure during the first year of the pandemic, was successfully visualized using PCA and msPHATE, and showed global and more fine-scale structure for each wave. The fine level of msPHATE embeddings were able to identify an outlier cluster during the first wave that dominated during the second wave (haplotype XIV, Figure 5). Indeed, msPHATE embeddings can identify sub-lineages with the potential of becoming a variant of concern, and can recapitulate hierarchical relationships between sub-lineages.

In conclusion, we presented a series of computational and population genetics tools to ease the lineage tracing of SARS-CoV-2 variants and predict emerging ones. We also presented a novel application for msPHATE as a method to analyze viral genetics data, and as a lineage surveillance method. Together, these approaches constitute a comprehensive toolbox to allow the scientific community to continually and closely monitor the evolution of any viral population. In the case of the ongoing pandemic caused by the SARS-CoV-2 virus, developing a dynamic, global and up-to-date understanding of viral evolutionary strategies will be of utmost importance to rapidly respond to emerging variants, identify increasingly infectious or vaccine-resistant lineages, and identify at-risk populations.

## Methods

### Pre-processing details

We downloaded a total of 384,407 sequences from GISAID on January 19th 2021. We then removed samples from non-human hosts as well as those with incomplete sequences (< 29, 000bp), for a total of 339,427 sequences. Each consensus sequence was mapped separately to the SARS-CoV-2 reference genome (NC 045512.2) using minimap2 2.17-r974 (44). All mapped sequences are then merged back with all others in a single alignment bam file. The variant calling is done using bcftools mpileup v1.9. https://samtools.github.io/bcftools/ in haploid calling mode. Sequences are processed by batches of 1000 sequences to overcome technical issues in processing of very low frequency variants within a bam file. Once the variant calling is obtained for each batch, INDELS are removed and bcftools merge was used to merge all the variant calls across the entire dataset. Variants located in both ends of the genome, which have high-level of missingness (>20%, positions 1-54 and 29838-29903) were excluded. We then flagged spurious variants within these sequences (see below) and identified 361 samples with at least two flagged positions, which we removed from our dataset. Of the remaining 339,066 sequences, we excluded sequences without GISAID metadata and with incomplete sampling date (sampling month unavailable), which resulted in a final dataset of 329,854 high-quality consensus sequences. We divided this dataset into two pandemic waves: 139,515 sequences with a submission date between January 1st and July 31st 2020 were defined as first wave samples, and 190,339 sequences with a submission date between August 1st and December 31st 2020 where defined as second wave samples. A mutation database is built using the sqlite3 library. Only positions that are variant from the reference, including missing calls, were included in the *position* table of the SQL database. For phylogenetics analyses, the multiple sequence alignment of 360,026 consensus sequences from 2020 provided by GISAID was downloaded on May 12th 2021.

### Spurious sites flagging

Positions that were “masked” by De Maio et al. (45) were removed. We additionally developed a tool to flag spurious variant within consensus sequences due by sequence misalignment in original labs, which we initially detected by manually inspecting consensus sequences. These errors were found in larger proportion in the sequences uploaded to GISAID in the early stages of the pandemic. Our approach identifies substitutions compared to the reference genome that are located within 10 genomic positions of stretches of “N”, defined as at least 5 consecutive Ns. This strategy was applied to the 339,427 consensus sequences from human host, we identified 2,164 sequences with at least one flagged position (0.6%). Among these, flagged positions where the mutated allele otherwise reached 1% in one of the pandemic waves (for a total of 199 positions) were considered as real mutations and not as spurious sites. A total of 6,736 spurious sites were detected and the variant allele was replaced by N in the sequences. Furthermore, we removed 361 samples with at least two flags. Additional details on this procedure can be found in Supplementary Text. The code is available here https://github.com/HussinLab/covid19_mostefai2021_paper

### Imputation

For the 199 positions reaching 1% derived allele frequency (DAF) in the consensus sequences of one of the two pandemic waves, we imputed the missing alleles using ImputeCoVNet (16). Briefly, ImputeCoVNet is a 2D convolutional ResNet autoencoder that aims at learning and reconstructing SARS-CoV-2 sequences with the help of two sub-networks: (1) an encoder which is responsible to embed the given input into a low-dimensional vector, and (2) a decoder that is responsible for reconstructing that sequence from that low-dimensional vector. During training, the encoder network takes as input complete sequences encoded with a one-hot representation and the decoder outputs a reconstructed version. Once trained, the model is used to infer missing values within incomplete sequences: the missing alleles at previously defined positions of interest are predicted by the model during reconstruction. We evaluated imputation accuracy on sequences without missing data and reached an accuracy of 99.12% on this set of mutated positions. The code is available here https://github.com/HussinLab/covid19_mostefai2021_paper

### Identification of high-frequency representative substitutions

The 329,854 high-quality consensus sequences were binned into months according to collection date. Monthly DAF for each substitution (24,802) was computed using consensus sequences available for each month. A total of 53 substitutions with DAF over 10% in at least one month were considered as high-frequency substitutions, 20 in the first wave and 33 in the second wave. In the second wave, the DAF trajectories (ie. DAF per month for each mutation) were highly correlated (Pearson *r*^2^ > 99%), forming two distinct groups of substitutions: for each of the groups, only one mutation was retained, genomic positions 22227 and 23063. With the 20 high-frequency substitutions from the first wave, a total of 22 high-frequency substitutions were considered to be representative of the evolutionary trajectories of SARS-CoV-2 in 2020 (Table S2).

### Haplotype network

The 22 representative positions were used to define viral haplotypes, which consist of group of alleles at these positions that are inherited together from a single parental sequence. We obtained a total of 463 unique haplotypes, and only those with more than 50 sequences were kept, for a total of 56 haplotypes. A haplotype network representing distance relationships between haplotypes was built. Because SARS-CoV-2 sequences were sampled sequentially through time, the haplotype network takes the temporal dimension into account. We split the year into 24 intervals representing half-months and each consensus sequence was attributed to one time interval. For each time interval, a haplotype network is generated using the haplonet function of the pegas R package (46) by including only sequences that occurred before or within this time interval. The networks were merged iteratively over time. At each step, if the merging creates a cycle (ie. the addition of a haplotype that is linked with two previously linked haplotypes) we removed the branches of the cycle that link the haplotypes for which the first occurrence was the longest timeframe. If many links had the longest timeframe, we removed the link between the more differentiated haplotypes. This process solved several time inconsistencies. The code is available here https://github.com/HussinLab/covid19_mostefai2021_paper

### Phylogenetic analysis and molecular clock

To reduce the dataset to allow feasible phylogenetic analyses, we applied several filters. (1) We kept only sequences from the 17 main haplotypes were kept and identical consensus sequences were merged, keeping the earliest collection date as the annotation. (2) Outlier sequences in term of their number of mutations at a given date were excluded. (3) We sampled at least 3 samples per date per haplotype and then balanced the sampling up to a maximum of 1000 samples per haplotype. The resulting dataset has a total of 15,690 sequences. The sites identified as problematic for phylogenetic tree reconstruction (problematic sites list v. 2021-04-15) were removed (45). The phylogenetic tree is computed using FastTree v2.1.11 (47) using a GTR + Gamma model. The divergence tree is then refined using TreeTime (v. 0.7.4) (48; 49). The root-to-tip distance was computed using TempEst v1.5.3 (50) and tree visualization was made using ggtree (51). Further improvement can be done to the alignment by removing poorly aligned regions using Gblocks program (52). However, this approach removes actual real positions from the alignment and make the tree less precise within sub-lineages. The code is available at https://github.com/HussinLab/covid19_mostefai2021_paper

### Tajima’s D

To track SARS-CoV-2 haplotypes’ spread, we used a population genetic metrics that can infer changes in effective population sizes by comparing the contribution of low- and intermediate-frequency mutations to viral genomic diversity, i.e. Tajima’s D (Tajima 1989). We calculated Tajima’s D at the continent level to be able to relate its time series to the haplotype network. For each haplotype and each of the 12 months of the first and second waves, we randomly sampled 20 viral consensus sequences from each continent to calculate Tajima’s D, and repeat this procedure 500 times to obtain confidence intervals. Lineages or time bins with fewer than 20 sequences were discarded. This sub-sampling method allowed us to control for differences in sample size across continents and time, although sampling biases inevitably result in reduced detected diversity. After calculating Tajima’s D, we correlated it to the number of sequences per haplotype per continent per month, which we used as a proxy for the number of cases per haplotype per continent per month. We also performed this correlation for each haplotype separately. We evaluated the significance of the correlation using the permutational ANOVA (n=5000 permutations) implemented in the R package “lmPerm” (v.2.1.0) (53). These analyses were implemented in R and are available on Github (https://github.com/arnaud00013/SARS_CoV_2_haplotypes_Tajima_D_2020_time_series/).

### Dimensionality reduction techniques

Principal Component Analysis (PCA) was performed on the first and the second waves high-quality consensus sequences. Genomic positions with more than 10% of missing samples were removed from analysis: 247 for the first wave and 6 for the second. We only kept the each derived alleles at a position when seen in at least 10 samples, which resulted in a final set of 6163 mutated positions for the first wave and 9818 for the second one. Triallelic and Quadriallelic sites are here coded as separate mutations. Missing data is encoded as reference allele. We used the incremental PCA method (54). Uniform Manifold Approximation and Projection (UMAP) is a python library that was also used on the data projected on the 20 first components of the PCA using the default parameters of the algorithm. In order to have the better fit, we explore each pair of those parameters: minimum distance [0.01, 0.05, 0.3, 0.8, 0.99] and neighbor numbe:r [10, 50, 80, 100, 200]. The code is available at https://github.com/HussinLab/covid19_mostefai2021_paper. Finally, msPHATE (13) (package available at https://github.com/KrishnaswamyLab/Multiscale_PHATE) embeddings were computed for the first and second waves GISAID high quality consensus sequences. A 0123 encoded matrix was used for each wave to generate the MS-PHATE embeddings. For a given position, the 0123 encoding is as follow: 0 encodes for the reference allele, and 1 2 and 3 encode for the 3 derived allele in decreasing frequency order independently for each wave. MS-PHATE returns embeddings and clusters at all granularities and we chose salient levels based on gradient analysis for visualization. Each dot in the plot consists of a group of samples in the original data, and we color the dots using the majority vote of haplotype. MS-PHATE provides the relationship between groups existing at different granularities and we use that to identify subgroups on finer levels comparing to a coarse level.

## Data and code availability

The code is available here https://github.com/HussinLab/covid19_mostefai2021_paper, the data is hosted on GISAID (1)

## Acknowledgments

This study was supported by funding from the Fonds de recherche du Québec – Santé (FRQS), Canada Foundation for Innovation, IVADO COVID19 Rapid Response grant (CVD19-030), the Montreal Heart Institute Foundation, the National Sciences and Engineering Research Council (NSERC), Alliance COVID-19 Grant (#ALLRP 554923–20), and the Canadian Institutes of Health Research (CIHR) (#174924), National Science Foundation (NSF: # 1636933 and # 1920920). E.C. and J.H. are FRQS Junior 1 Research Scholars. This work was completed thanks to computational resources provided by Compute Canada clusters Graham and Beluga.

## Supplementary Material

### Supplementary Text

#### Additional Details on Phylogenetic Analyses

A multiple sequence alignment of 360,026 consensus sequences provided by GISAID was downloaded (version of January 20th 2021, downloaded on May 12th 2021). Sequences annotated by the 17 most frequent haplotypes were kept for further analysis. To avoid over-representation, identical consensus sequences were merged represented by the first one seen (earliest date). Only GISAID sequences with a date between January and December 2020 were kept. Samples diverging too much in regards to their sampling date were then removed following this equation j * 0,0597 + 50 where j is the number of days since the January 1st 2020. It means that we allow the mutation rate to be a bit above one mutation every 16 days. The factor 0,0597 was computed by making a linear regression between date and unique number of mutation per day.

Next, to obtain a result representative of the whole tree, we sampled at least 3 samples per date per haplotype and then completed the sampling up to a maximum of 1000 samples per haplotype. Sequences were put into a single fasta file, making sure we also have the presence of the Wuhan reference sequence representative (EPI ISL 402119 representing EPI ISL 402124, since the distance is null and the sampling date is older). The sites identified as problematic for phylogenetic tree reconstruction (flagged as “mask” in the problematic sites list v. 2021-04-15)[ref] were removed from the alignment using the relative reference genome positions on the multiple alignment. The phylogenetic tree is then computed using FastTree v2.1.11 (47) using a GTR + Gamma model. The divergence tree is then refined using TreeTime (v. 0.7.4) (48). Without any corrections, the constructed phylogenetic tree using FastTree splits haplotype I (Figure 2C), and combined very distinct lineages haplotype I and II. Because of our haplotype network, we were able identify these connection issues and manually correct them. By removing problematic connections and sequences, we obtained a more representative phylogenetic tree with the correct connections and successions of SARS-CoV-2 lineages during the first year of the pandemic (Figure S3). To measure the mutational rate of each haplotype through the pandemic’s first year we first used a root-to-tip molecular clock approach (Figure S5A) that was run on the divergence tree (Figure 2C). The root-to-tip distance was calculated using TempEst v1.5.3 (50) and tree visualization was made using ggtree (51). One sample was removed from the tree and the root-to-tip graph (EPI ISL 833364) due to a incorrect sampling date (which is January 8 2020, prior to SARS-CoV-2 genome sequencing started), resulting in a total of 15,690 sequences.

This root-to-tip molecular clock approach is slow and thus needs heavy down-sampling of sequences. Therefore, for the same sequences, we calculated the number of mutations from the reference and plotted them along their date of sampling (Figure S5B). This latter approach is faster and takes advantage of all sequences available for a given time. Additionally, with this approach we can see some mutational “jumps” that are less apparent with the molecular clock. For example, in February, we see some haplotype VII samples with a mutational jump (Figure S5B) that were as evident using the molecular clock (Figure S5A).

#### Mutational landscape of msPHATE clusters II and III

The finer level representation of the first wave sequence data reveals three separate clusters of haplotype II, and one of haplotype III clusters splits into five sub-clusters (Figure S9B) and each subgroup mutational landscape can be inspected. The mutational graphs show that each haplotype II MS-PHATE cluster has specific mutations defining it, in addition to mutations defining haplotype II (Figure S9). Cluster 3 (C3) is defined by the lack of additional mutations suggesting this sub-lineage as the ancestral; Cluster 2 (C2) is defined by three mutations (ORF1a:S3884, NSP14:A6250V, and an intergenic (IG) mutation); However, Cluster 3 (C3) does not seem to have a particular mutational signature (suggests that a finer level will split this group into multiple sub-lineages).

The mutational graphs show that each haplotype III MS-PHATE cluster has specific mutations defining them, in addition to mutations defining haplotype III (Figure S9). Clusters 1 (C1), 2 (C2) and 3 (C3) do not seem to have a specific mutations defining them, suggesting that a finer level could reveal more sublineages; Clusters 4 (C4) and 5 (C5) are respectively defined by two mutations at genomic positions 20268 and 29734, and one mutation at genomic position 15324. However, cluster C6 corresponds to haplotype XIV which became a dominating lineage in GISAID consensus sequences during during the second wave of the pandemic.

#### Details on Spurious Sites Flagging

Upon manual inspection of consensus sequences, we noticed a systematic effect impacting regions of the genome around stretches of N for sequences from specific sequencing centers. We thus developed a script to evaluate this effect automatically, that identifies and removes spurious mutations resulting from this type of sequencing and assembly errors.

Our script takes as input FASTA file of sequence alignment (based on the reference genome) and the fasta file of the reference sequence. The sequences in the alignment must be exactly the length of the reference genome (here, 29 903 nucleotides). Two parameters can be specified: the first parameter (alpha) corresponds to the minimum number of consecutive unknown nucleotides (N) in a stretch (see Figure 2); the second parameter (beta) corresponds to the maximum distance between a putative variant and the stretch of Ns in number of nucleotides. Here, we used alpha=5 and beta=10.

In a specific sequence from the FASTA alignment file, the script will find all stretches of at least alpha consecutive Ns. Each nucleotide within beta positions of the every stretch of N will be compared to the nucleotide at the corresponding position in the reference genome: a position which differs will be identified as a putative spurious mutation (flagged) and replaced by an unknown nucleotide in the fasta file in the outputs. It is also possible to specify a bed file of positions to investigate. In this case, the script won’t look at all position, but rather will only flag positions specified in the bed file. This option speeds up processing if the user is only interested in a specific region of the genome. We also added an option to protect specific positions from being flagged, such as known real mutated positions that could inadvertently be located next to stretches of Ns. The user can give a bed file with or without a column containing the alternative allele at the protected position. If there is no alternative allele specified, the position will not be considered as spurious (but will be reported). If an allele is specified, the script will consider it as spurious only if the nucleotide at that position differs from the reference and from the specified alternative allele.

The output the script consists of three files: a corrected alignment file in FASTA format where flagged positions are replaced by Ns; a file reporting the number of flagged sites per sequence; and a file with information on every flagged site: sequence ID, position, the distance from the closest stretch of Ns, the flag, and the nucleotide found at this position. The flag is set to 0 when the site is spurious, set to 1 if the site is protected, or set to 2 if the site is protected but differs from the alternative allele reported. Only positions flagged with 0 and 2 are replaced by N in the output FASTA file.

#### Time-dependent Haplotype Network Optimization

For each consensus sequence, the sampling date is rounded into half-month date intervals. Next, each haplotype is associated to a date interval by identifying its first occurrence. For each interval, a network was generated using pegas (46) with only haplotypes that occurred before or within the interval. The networks were then merged iteratively over time. To avoid circular connections between haplotypes during the merging process, the circular patterns in the network are removed by first identifying these patterns then removing the links between networks with the longest time interval difference. However, if the merging creates a cycle, we only keep the branches of a cycle that link the haplotypes with the lowest time difference.

---

**Table S1.** Mutation Frequency Spectrum. Number of mutated genomic positions within the GISAID sequences binned by the whole year, the first, or the second wave of the pandemic. The rows represent the mutated stratified by their appear in each group of sequences.

| Number of<br>consensus<br>sequences with<br>mutation | Total 2020<br>mutation counts | First Wave<br>mutation counts | Second Wave<br>mutation counts |
| --- | --- | --- | --- |
| 1 | 3,168 | 4,758 | 3,695 |
| 2 - 10 | 9,877 | 9,776 | 9,471 |
| 11 - 99 | 8,622 | 4,862 | 6,646 |
| >100 | 3,135 | 1,007 | 2,398 |
| total | 24,802 | 20,403 | 22,210 |

**Table S2.** The 22 mutated genomic positions that represent the genetic diversity of SARS-CoV-2 during the first year of the pandemic. These 22 positions define the haplotype groupings. For each mutetd position, the frequencies per wave and for the whole year are also represented. Additionally, for each position, we report their substitution, gene position, amino acid change, and functional consequence on the resulting protein.

| Genomic<br>position | Frequency<br>2020 | Frequency<br>first wave | Frequency<br>second wave | Nuc acid<br>Substitution | Gene | Amino acid<br>Substitution | Functional<br>consequence |
| --- | --- | --- | --- | --- | --- | --- | --- |
| 241 | 94% | 88% | 100% | C>T | intergenic | N/A | N/A |
| 313 | 4% | 7% | 1% | C>T | nsp1 | L16L | synonymous |
| 1059 | 14% | 21% | 8% | C>T | nsp2 | T265I | missense |
| 1163 | 5% | 7% | 2% | A>T | nsp2 | I300F | missense |
| 3037 | 94% | 88% | 100% | C>T | nsp3 | F924F | synonymous |
| 7540 | 3% | 6% | 0% | T>C | nsp3 | T2425T | synonymous |
| 8782 | 2% | 5% | 0% | C>T | nsp4 | S2839S | synonymous |
| 14408 | 94% | 88% | 100% | C>T | nsp12 | P4720L | missense |
| 14805 | 3% | 4% | 2% | C>T | nsp12 | Y4852Y | synonymous |
| 16647 | 3% | 6% | 1% | G>T | nsp13 | T5466T | synonymous |
| 18555 | 3% | 6% | 1% | C>T | nsp14 | D6102D | synonymous |
| 22227 | 23% | 1% | 44% | C>T | S | A222V | missense |
| 22992 | 6% | 6% | 6% | G>A | S | S477N | missense |
| 23063 | 5% | 0% | 10% | A>T | S | N501Y | missense |
| 23401 | 3% | 6% | 0% | G>A | S | Q613H | missense |
| 23403 | 94% | 88% | 100% | A>G | S | D614G | missense |
| 25563 | 22% | 27% | 17% | G>T | ORF3a | Q57H | missense |
| 26144 | 2% | 3% | 0% | G>T | ORF3a | G251V | missense |
| 28144 | 2% | 5% | 0% | T>C | ORF8 | L84S | missense |
| 28881 | 34% | 41% | 28% | G>A | N/orf9c | R203K | missense |
| 28882 | 34% | 41% | 27% | G>A | N/orf9c | N203N | synonymous |
| 28883 | 34% | 41% | 27% | G>C | N/orf9c | G204R | missense |

**Fig. S1:**
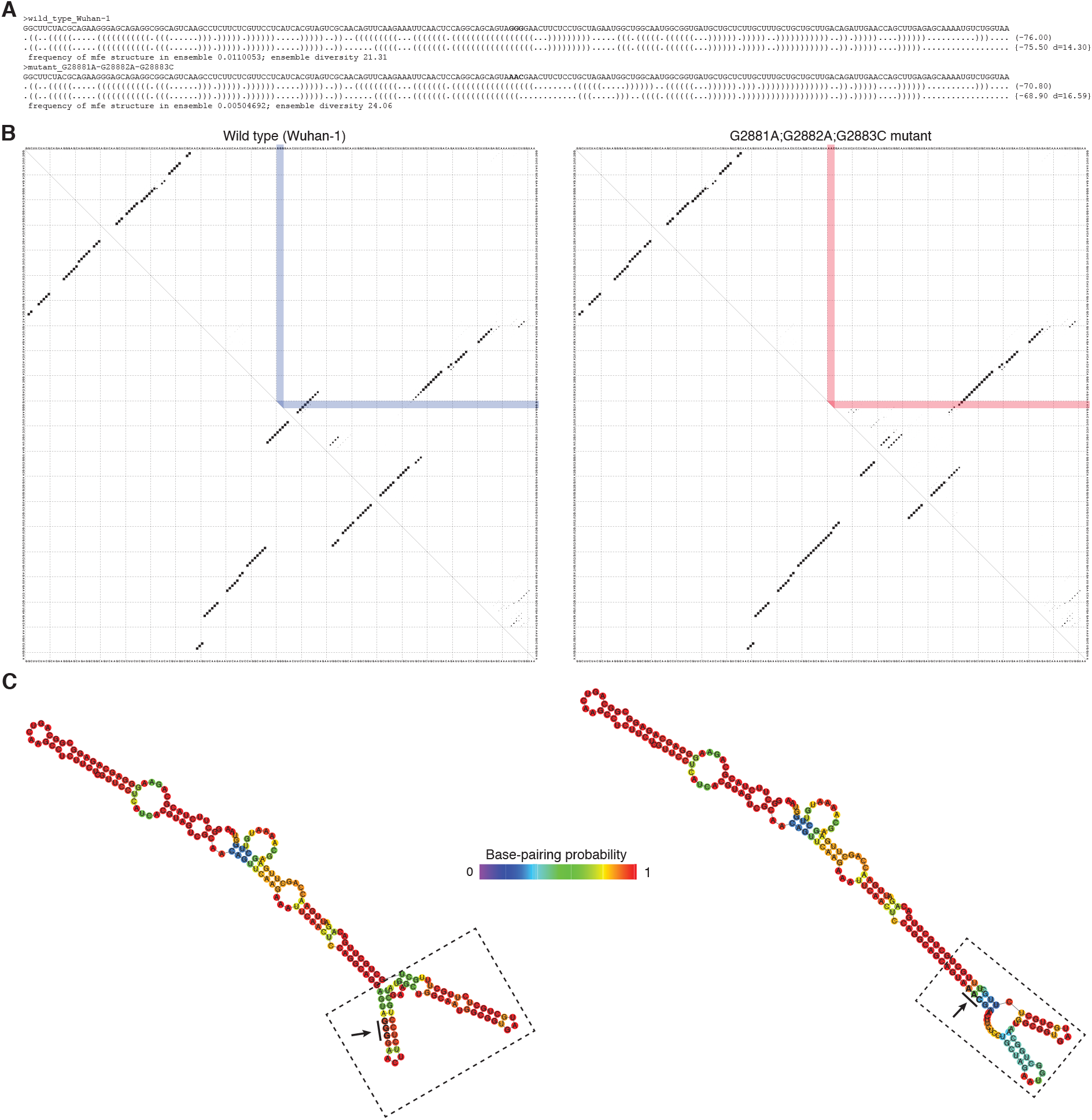
Comparative secondary structure prediction of reference SARS-CoV-2 transcript *vs* trinucleotide ORF9c mutant. (A) Primary sequence, minimum free energy (MFE) RNA secondary structure prediction (RNAfold with partition function, ViennaRNA package) and centroid structure prediction with MFE frequency in – and diversity of – Boltzmann ensemble. (B) Base-pair probability matrices for reference sequence (left) and trinucleotide mutant sequence (right) with the position of affected nucleotides highlighted in color. The size of dots reflects the base pairing probability distribution in the Boltzmann ensemble of suboptimal base pairings (right of diagonal), whilst the MFE prediction is illustrated left of the diagonal. (C) RNA secondary structure predictions annotated with base-pairing probabilities. Dotted boxes highlight the topological differences associated with the trinucleotide mutation (arrow).

**Fig. S2:**
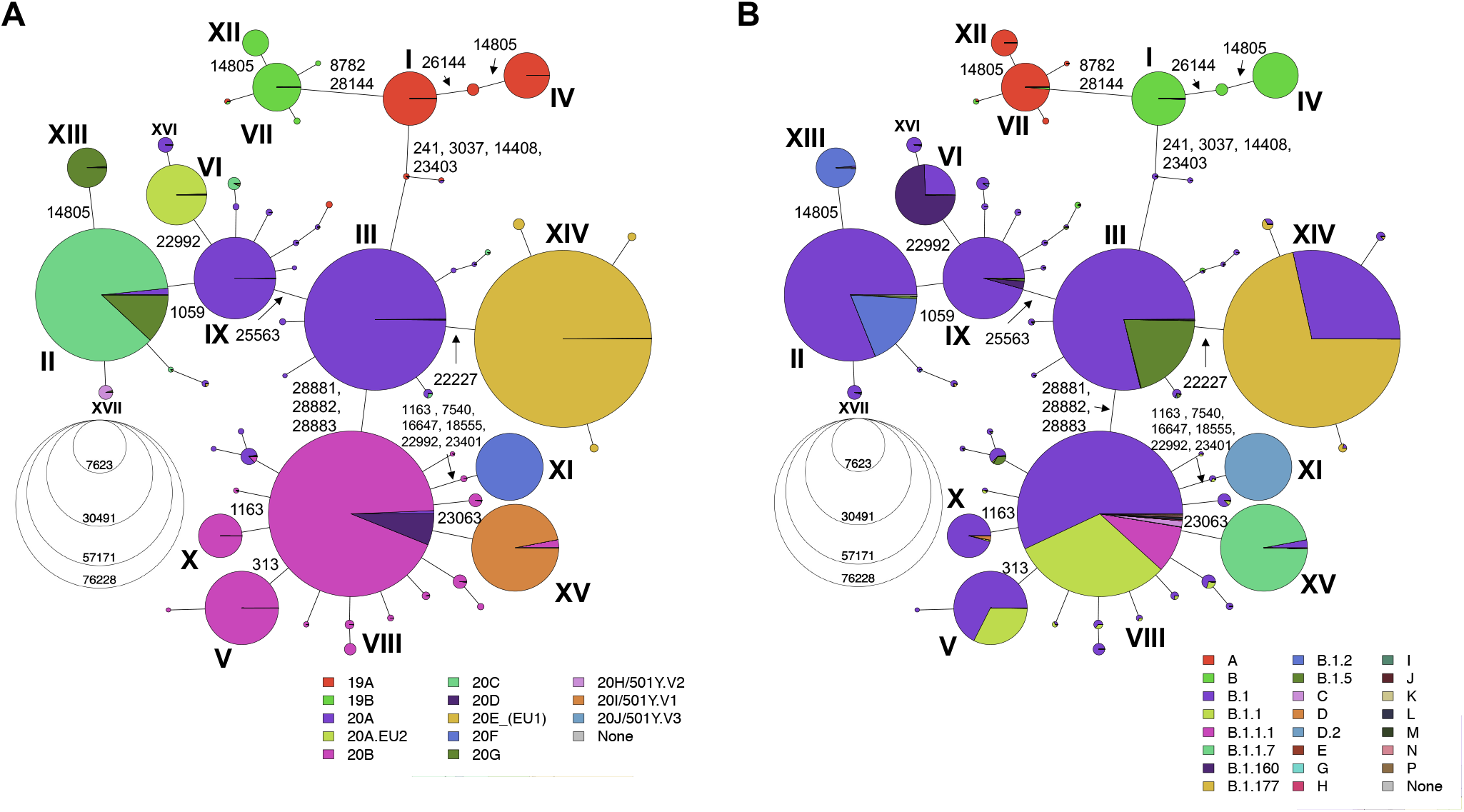
Haplotype Networks with NextStrain and Pangolin lineage correspondence. (A) Haplotype Network with NextStrain correspondence. Each branch represents the mutation that define the haplotype lineage. (B) Haplotype Network with Pangolin lineage correspondence. The pangolin lineage has a color only if there is more than 5000 samples with this lineage. Those with at most 5000, we added them to the sub-lineage (the letter+one number) and if even this sub-lineage is no more than 5000, we only kept the first letter.

**Fig. S3:**
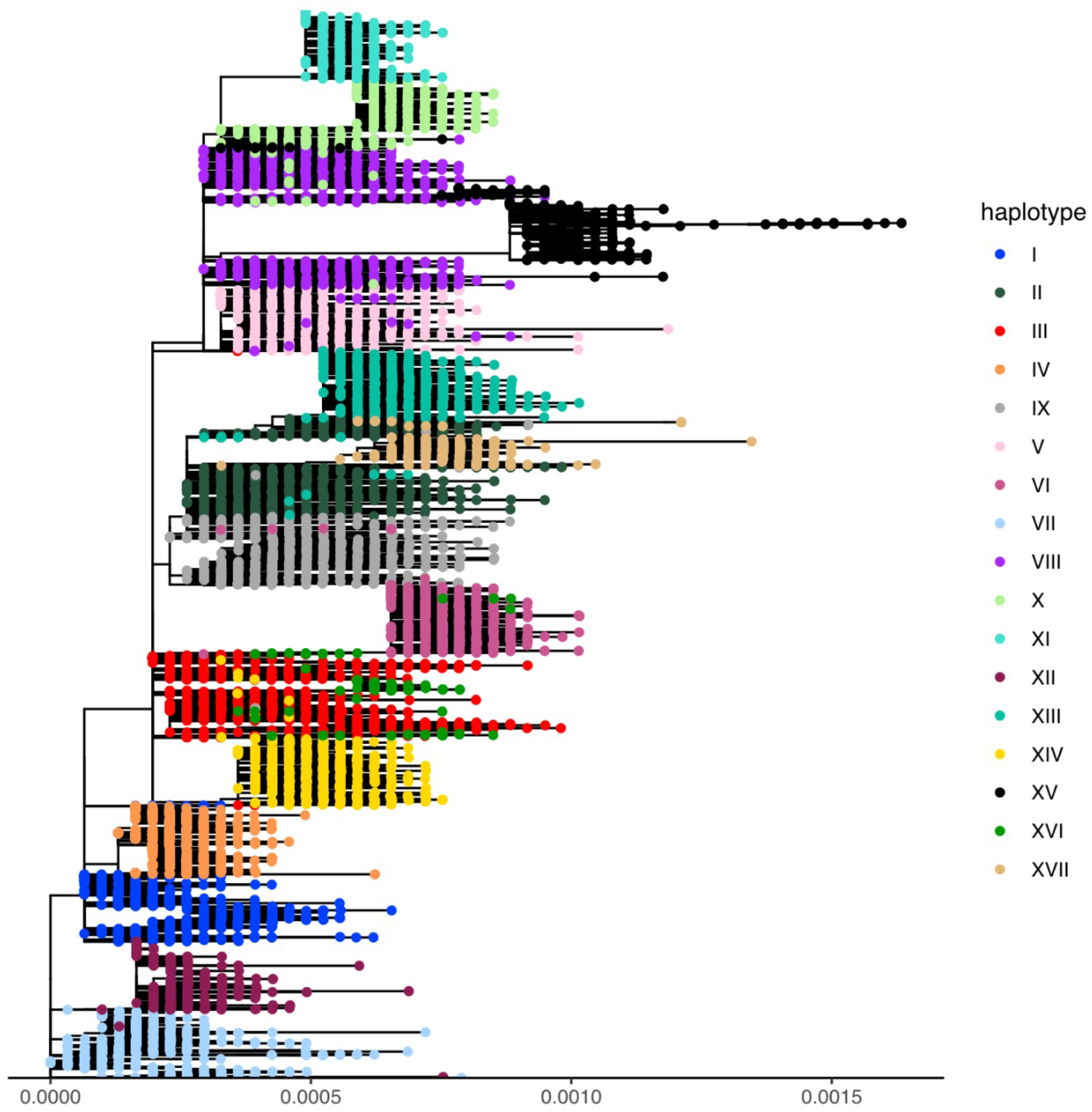
Phylogenetic tree summarizing the circulating lineages during the first year of the pandemic. The phylogenetic tree was built using the same samples as what is seen in Figure 2C, the alignment was post-treated using the program Gblocks (52), which eliminates poorly aligned positions and divergent regions of a multiple sequence alignment. Default parameters were used. The tree is then built using FastTree and TreeTime using a GTR+Gamma model.

**Fig. S4:**
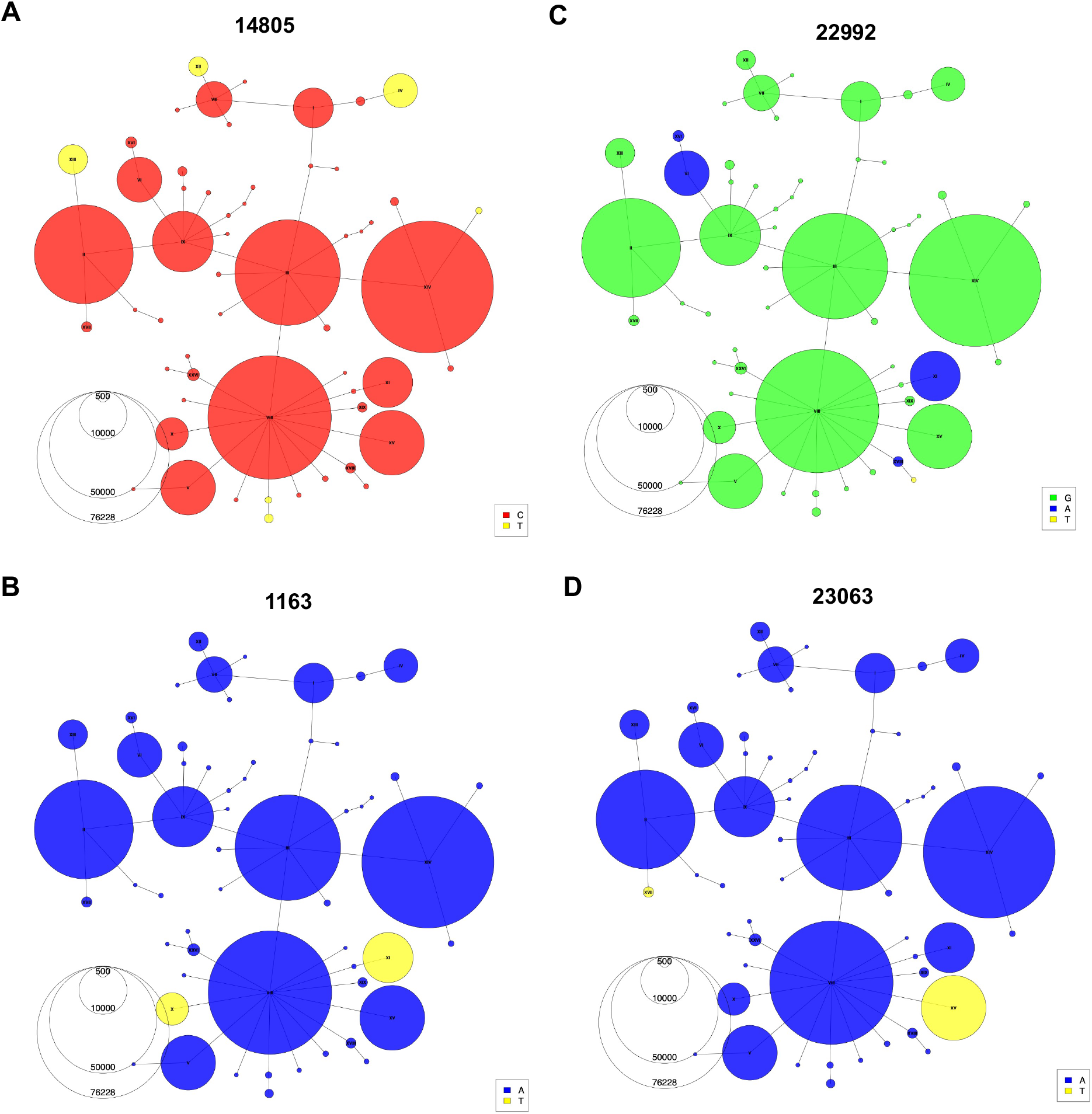
Recurrent mutations visualized using the haplotype network. Haplotype Networks colored according to the presence of specific alleles at genomic positions 14805 (A), 1163 (B), 22992 (C), and 23063 (D).

**Fig. S5:**
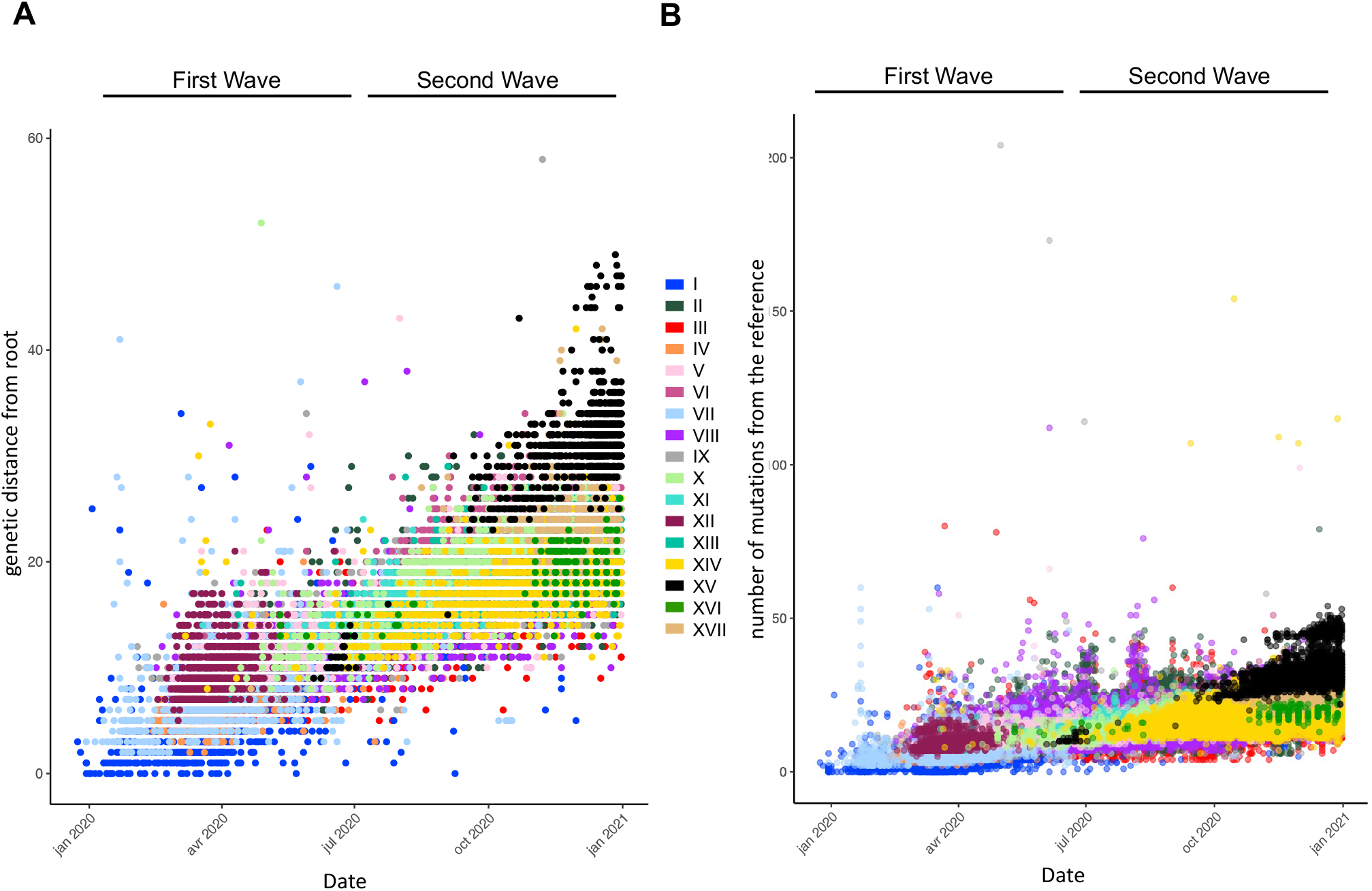
Different approaches to measure mutational jumps in lineages. (A) Root-to-tip distance plot based on divergence phylogenetic tree of GIAID SARS-CoV-2 consensus sequences. The root-to-tip distance is measured from the phylogenetic tree presented on Figure 2C using TempEst v.1.5.3 (50). Then labeled using haplotype annotations. (B) Number of mutations compared to the reference genome (NC 045512.2) for the same samples as the ones in panel A and labeled using the haplotype annotation.

**Fig. S6:**
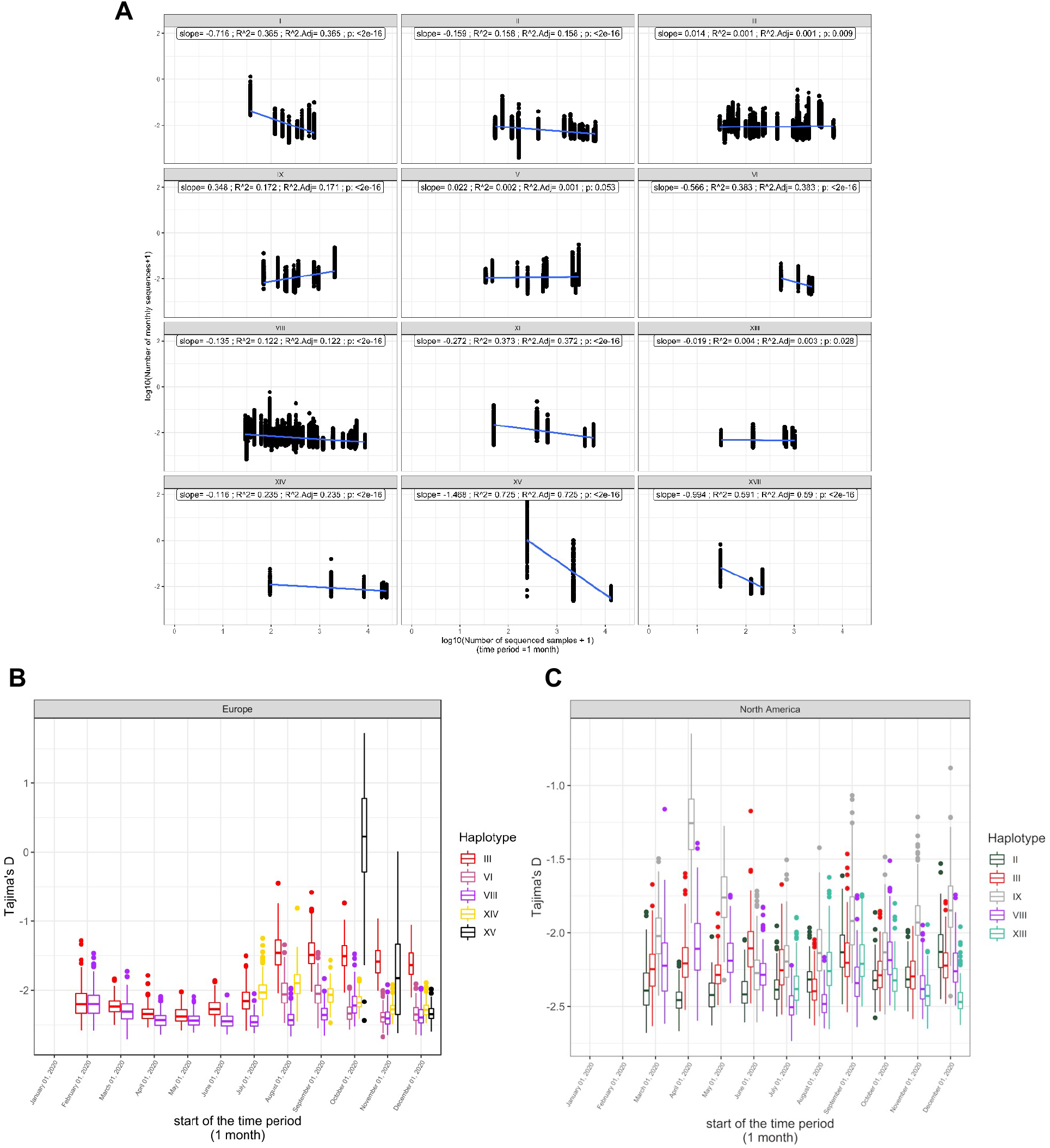
Tajima’s D correlation with haplotype trajectories over time in each continent. (A) *R*^2^ of Tajima’s D across haplotypes. For most of the haplotypes, Tajima’s D correlates negatively and exponentially with the number of sequences per haplotype per continent per month. This value is used as a proxy for the number of cases per haplotype per month per continent. For almost all the haplotypes (I, II, III, IX, VI, VIII, XI, XIII, XIV, XV, and XVII), the correlation is significant (PERMANOVA p ¡ 0.05) with a mean adjusted R2 of 0.24 across haplotypes (s.d.=0.23). This correlation is consistent with the fact that smaller negative Tajima’s D values are associated with stronger spread and that Tajima’s D can be used for phylodynamic inferences. The moderate correlation strength (R2) can be explained by the fact that the number of sequences is probably an underestimate of the number of true cases and by the fact that the analysis captures the average trend in each continent which is weakened by variations of epidemiological dynamics across countries and cities. Tajima’s D estimates of the five most prevalent haplotypes in Europe (B) and North America (C) for the first year of the pandemic. Boxplots represent 500 resampled estimates of Tajima’s D from random resampling of 20 genome sequences for each month with at least 20 sequences.

**Fig. S7:**
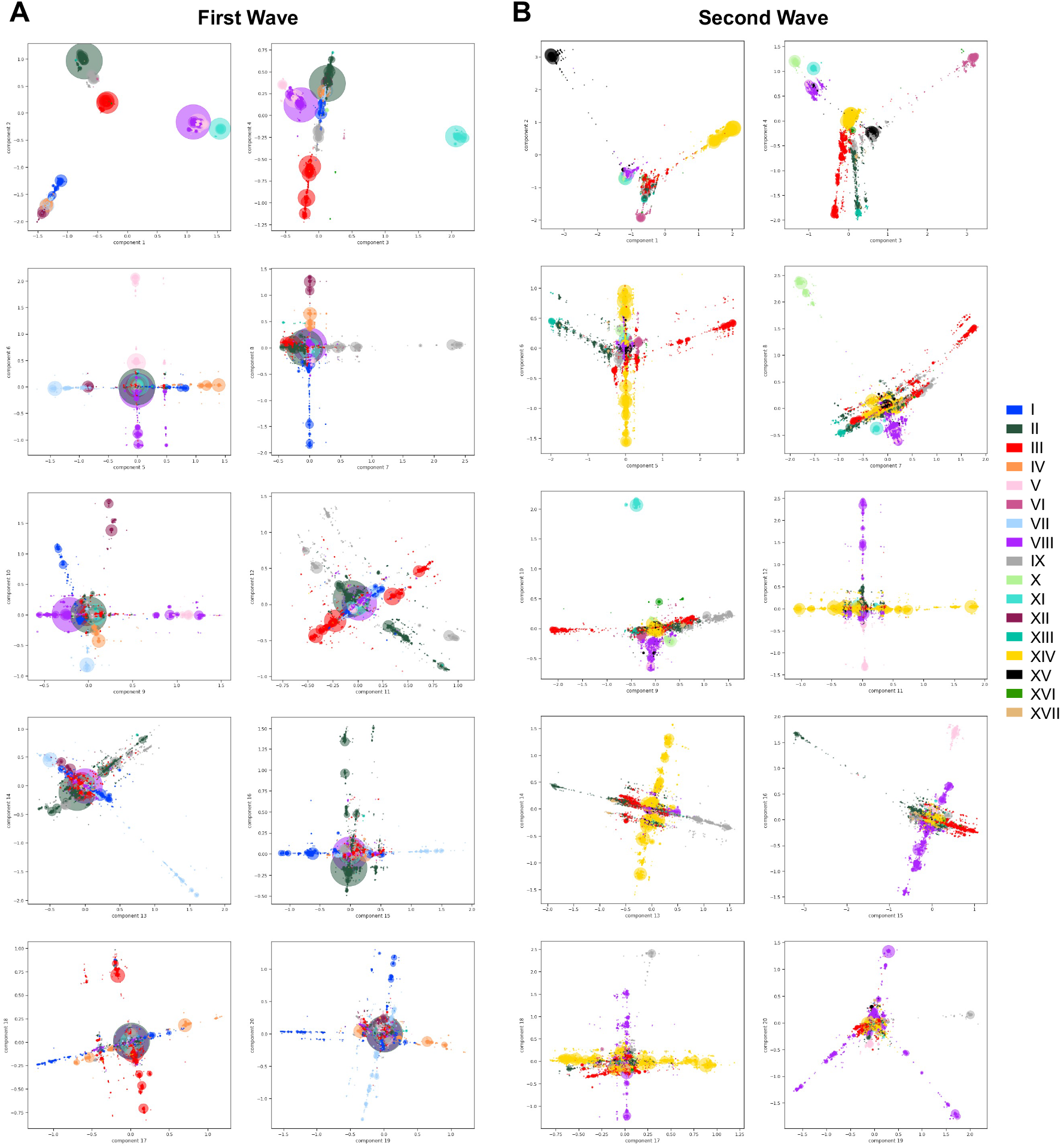
Viral population structure during the first and second wave of the pandemic across all 20 Principal Components (PCs). Principal Component Analyses (PCA) of genetic diversity of the first (A) and second (B) wave consensus sequences shown for the 20 first PCs. Genetic variation present in at least 10 genomes is used. The PCA is computed with all sequences, and only the sequences from the 17 main haplotypes are projected. Identical sequences are projected onto the same coordinates, therefore the number of sequences represented by each point is proportional to the size of the dots, with added transparency.

**Fig. S8:**
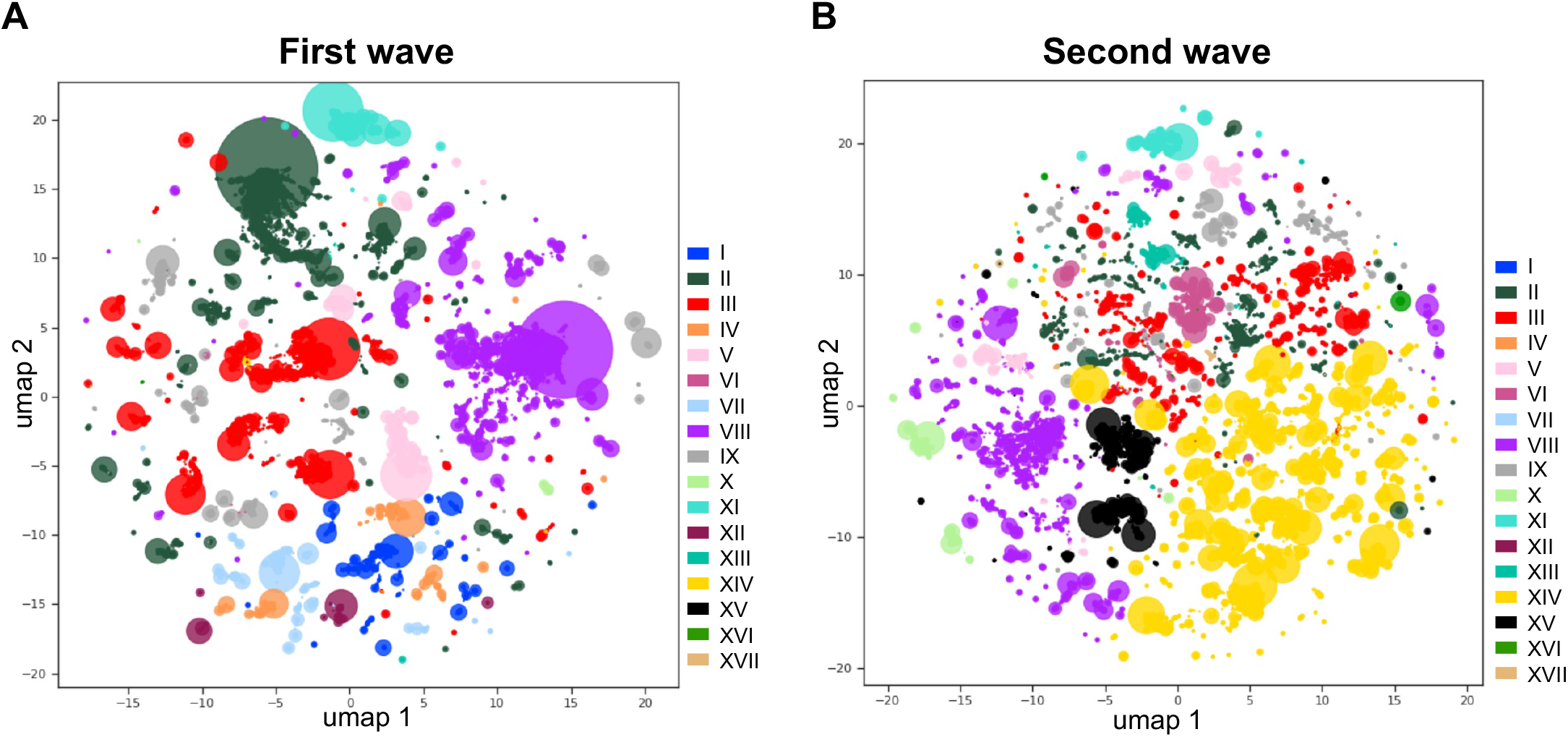
Viral population structure during the first and second wave of the pandemic visualized with 2-dimensional Uniform Manifold Approximation and Projection (UMAP). UMAP performed on the 20 principal components (PCs) generated from genetic variation found in GISAID consensus sequences of the first (A) and second wave (B) of the pandemic (Figure S7), colored according to the 17 main haplotypes. To generate this UMAP embedding, the following parameters were used: *n_neighbor* = 50, *min_dist* = 0.3.

**Fig. S9:**
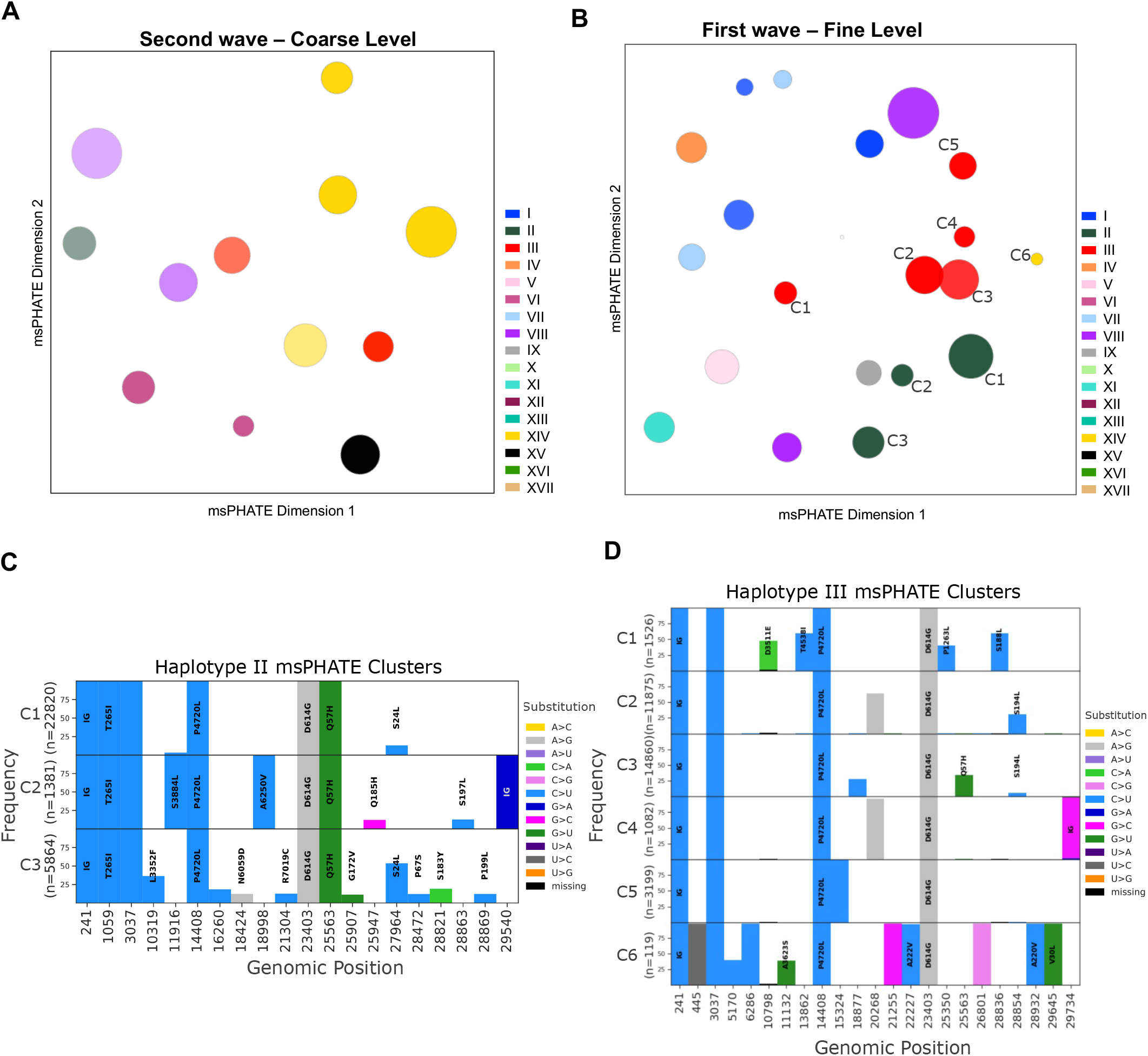
Investigation of msPHATE viral stratification. (A) Corse grain visualization of msPHATE embedding generated from genetic diversity of the second wave consensus sequences. The clusters are colored by majority vote. (B) Annotated fine grain visualization of msPHATE embeddings of the genetic diversity of the first wave as seen in figure 5B. Mutated positions in each cluster group of haplotypes II (C) and III (D) annotated in B are listed along the x-axis in the order they appear on the genome. Mutations in at least 25% of the samples of each cluster are represented as a bar, and the corresponding amino acid changes are shown on the bars (blank bars represent synonymous mutations).

**Fig. S10:**
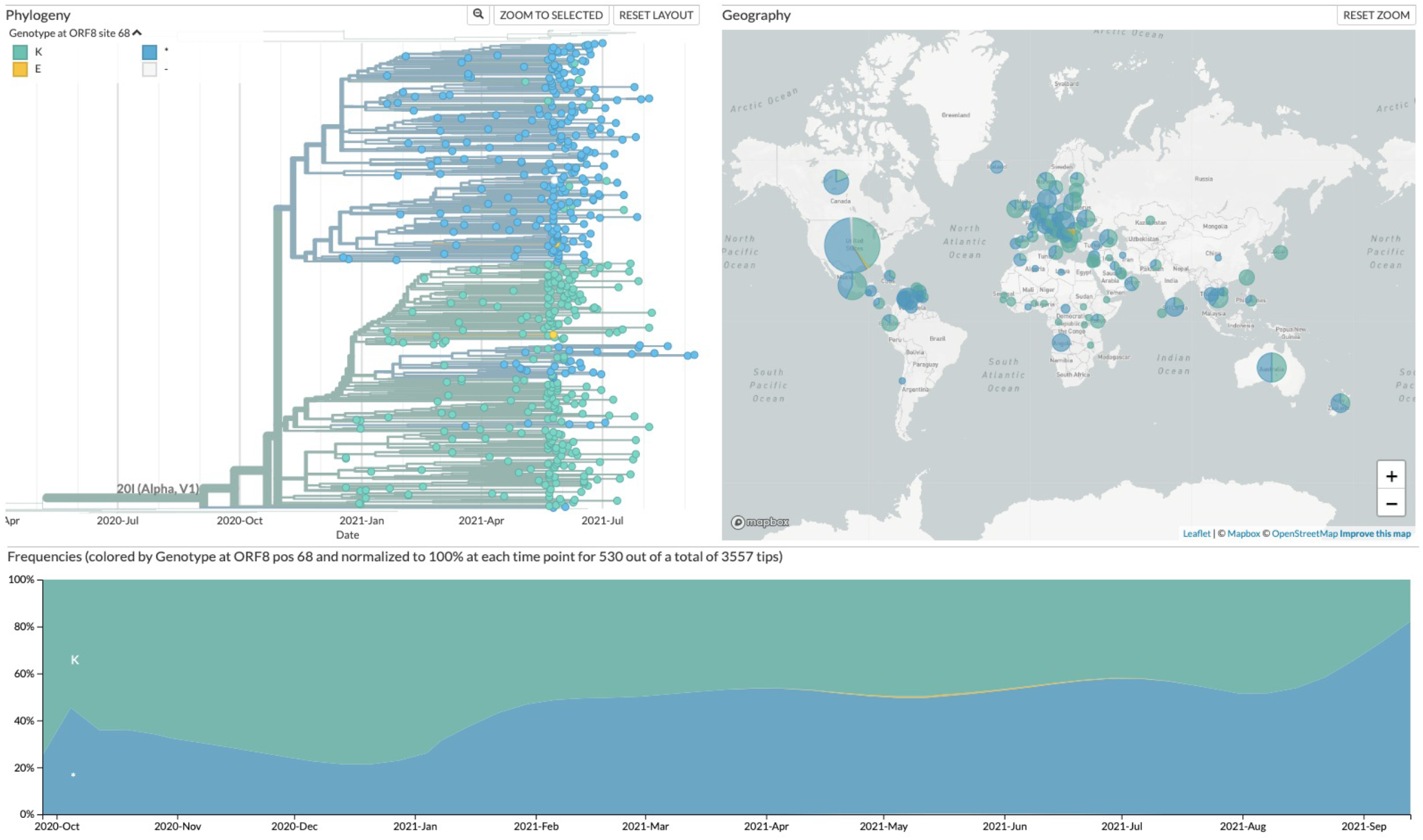
NextStrain frequencies of mutation A29095U (ORF8:K68*) on the Alpha variant identified by msPHATE. NextStain phylgentic tree, word-wide frequency, and time-series frequency of samples with mutation ORF8:K68*. Figures captured from nextstrain.org/sars-cov-2 on September 29th 2021.

